# Tricellular junction recruitment of the Wave regulatory complex by Sidekick and Lar induces protrusive activity resolving cell intercalation

**DOI:** 10.1101/2024.06.28.599016

**Authors:** Lisa Calvary, Hervé Alégot, Pierre Pouchin, Graziella Richard, Caroline Vachias, Vincent Mirouse

## Abstract

Cell intercalation, a fundamental morphogenetic process characterized by the exchange of neighboring cells, plays a pivotal role in epithelial tissue development. While the initiation of new junctions remains poorly understood, recent research indicates the involvement of tricellular junction actors. In this study, we explore the contribution of the WAVE regulatory complex (WRC), a critical regulator of branched F-Actin generation, in tissue elongation and cell intercalation within the Drosophila ovarian follicular epithelium. WRC localizes at tricellular junctions, where it orchestrates the generation of highly dynamic protrusions emanating from one cell and extending between the bicellular junctions of neighboring cells. This protrusive activity is essential for the initiation of new junctions in cells located at the extremities of these junctions. Furthermore, our findings indicate that WRC recruitment at tricellular junctions is a redundant process, involving the cooperative action of the two transmembrane proteins Sidekick and Lar. Disruption of this recruitment impairs protrusive activity, cell intercalation resolution, and tissue elongation, thereby mechanistically bridging molecular, cellular and tissular scales. Consequently, this elucidates a critical mechanism underlying epithelial morphogenesis through actin polymerization at tricellular junctions.

## Introduction

How cells change their respective position during development to allow morphogenesis is a major question. In epithelial tissues such rearrangements are often materialized by cell intercalation, a process usually associated with tissue elongation in which cells exchange neighbors in the epithelium plan ^1–3^. Cell intercalation, also called T1 transition, is mechanically driven by an oriented disequilibrium of the balance between cortical tension, induced by Myosin II, and adhesion, mainly due to adherens junctions. This instability can be generated by intrinsic anisotropic modulation of Myosin II activity or by extrinsic forces applied to the cells. Schematically, intercalation can be divided into three successive steps (Fig 1A). First, the preexisting junction shrinks. Once disappeared, it leads to the formation of a specific figure called a 4-way vertex corresponding to the junction point between the different cells involved in the intercalation. In some cases, more than 4 cells can converge to the same point leading to the formation of a rosette. Finally, the 4-way vertex will resolve, and a new junction will form and lengthen between the intercalating cells. Of notice, this last step is the main contributor of tissue elongation ^4^. Whereas initial focus has been made on the mechanisms allowing pre-existing junction shrinkage ^5–7^, several studies have started to uncover mechanisms promoting new junction initiation and lengthening ^4,8–10^. It is not yet clear whether all these mechanisms are identical depending on the studied tissue. Nevertheless, a general outcome is that intercalation resolution and the subsequent new junction lengthening emerge from a bias in Myosin II recruitment, being stronger in the cells at the new junction extremities than along this junction. Of notice, the tricellular junction protein Sidekick (Sdk) has been identified in different tissues as participating in such a mechanism ^10–12^. However, its general requirement for morphogenesis could be questioned because *Drosophila sdk* null mutants are viable and fertile.

**Figure 1.**
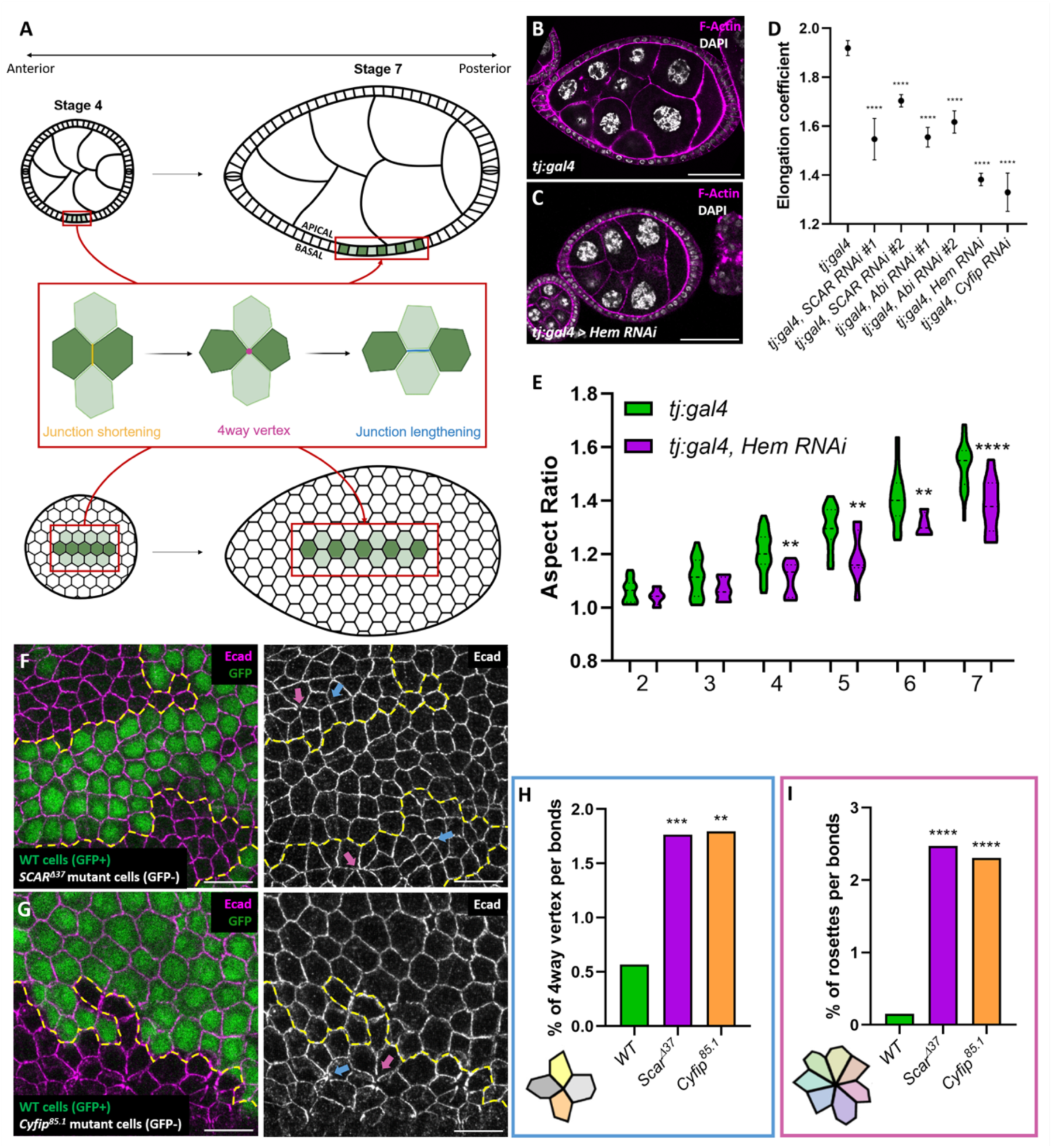
WRC is required for early elongation of follicles and cell intercalation resolution. A) Scheme of follicles observed in the middle plan (top) or at the surface (bottom) at stage 4 and 7. Middle plan shows the germ cell cyst surrounded by the follicular epithelium with its apical domain facing the germline. Surface view is in the plan of the epithelium. Zoom-in illustrates the intercalation process with junction shrinkage, formation of a 4-way vertex and intercalation resolution with the lengthening of a new junction. On all figures anterior is on the left and posterior on the right. B-C) Images of *tj:gal4* (control) (B) and *tj:gal 4, Hem RNAi* (C) follicles at stage 7. D) Elongation coefficient from stages 2 to 7 for the indicated genotypes. E) Measure of aspect ratio from stage 2 to 7 of control and *tj:gal 4, Hem RNAi* follicles F-G) follicles containing mutant clones for *SCAR* (F) and *Cyfip* (G) null mutations marked by the absence of GFP, and with an immunostaining against E-Cad. Blue arrows show 4-way vertices, pink arrows show rosettes. H-I) Quantification of the frequency of 4-way vertices (H) or rosettes (I). (Scale bars: 50μm on B,C, 10 μm on F,G)( ** P < 0.01, ***P < 0.001, **** P < 0.0001).

*Drosophila* ovarian follicle (also called egg chamber) provides a typical example of tissue elongation (Fig 1A). Follicles form initially a sphere composed of 16 germ cells, including the oocyte, surrounded by a monolayer epithelium of follicle cells. These follicles grow and differentiate through 14 developmental stages during which they elongate in their anteroposterior axis, prefigurating the shape of the fly embryos ^13^. This elongation is driven by the follicular epithelium ^14^. A first elongation phase relies on a double gradient of apical pulsations starting from the poles and creating tension along the anteroposterior axis ^15,16^. Consequently, cells become more constricted at the poles and more stretched and thinner at the equator. This first phase is also associated with cell rearrangements such as oriented cell intercalation and cell division ^15–17^.

Of note, this phase is independent of the Fat2 dependent planar cell polarity pathway that drives the second phase of follicle elongation starting around stage 8 ^15,18^.

Here, we identified branched actin and its control by the WAVE Regulatory Complex (WRC) as a critical actor of cell intercalation in follicle cells. Branched actin builds complex networks by the nucleation of new filaments on preexisting ones. This nucleation relies on the Arp2/3 complex that can be activated by different but related proteins or complexes, called Nucleation Promoting Factors (NPFs), including WRC ^19,20^. WRC is composed of 5 subunits: SCAR, Cyfip, Hem, Abi and HSPC300. WRC function is well described in migrating cells where it induces lamellipodia from the cell cortex. In neuro-epithelial cells of the fly retina, WRC activity has been implicated in junction lengthening, counteracting in a cell-autonomous manner myosin II activity (Del Signore et al., 2018). However, the role of WRC, and in a more general way of the actin cytoskeleton, during cell intercalation is still poorly explored.

We found that the WRC is required for the first phase of follicle elongation and cell intercalation resolution. WRC is found at tricellular junctions where it generates a specific subpopulation of F-actin that induces dynamics lamellipodia invading bicellular junctions of adjacent cells. This activity is required in the cells at new junction extremities for its lengthening. Finally, we deciphered how WRC is recruited at tricellular junctions. It involves direct interaction with Sdk protein, confirming recently published results ^21^. However, Sdk is not required to localize WRC at TCJs. We identified that the tyrosine phosphatase receptor Lar, which has been shown to recruit the WRC in another context, is also present at TCJs ^22,23^. Concomitant removal of both *sdk* and *Lar* is required to affect WRC localization at TCJs and phenocopies WRC loss on actin dynamics, cell intercalation and tissue elongation.

## Results

### WRC is required for follicle elongation and cell intercalation

In a reverse genetic screen to identify F-actin regulators involved in early elongation (stage 2-7), we identified the Wave Regulatory Complex (Fig 1B,C). Typically, knock-down (KD) using UAS:RNAi transgenes were induced with tj:Gal4, a driver specific for follicle cells. Elongation can be estimated by a coefficient that corresponds to the slope of the line given by the long axis in function of the short one, this method being independent of stage determination and allowing easy comparison between genotypes ^15^. Though a variability was observed, probably reflecting each RNAi line efficiency, many lines targeting different WRC subunits reproduced an early elongation defect, giving robust evidence of its involvement (Fig 1D). The elongation can be also represented using the aspect ratio (ratio between long and short axes) depending on the stage, showing that all the stages of early elongation were equally affected in WRC KD, a defect being detected as soon as stage 4 (Fig 1E). WRC KD blocks follicle rotation that is concomitant to early elongation, a process in which the whole follicular epithelium migrates around its anteroposterior axis and which is involved in egg elongation ^24,25^. However, rotation is not required for the early phase of follicle elongation but only for the subsequent one ^15,18^. Thus, WRC has at least one additional function promoting follicle elongation during these early stages.

We have previously shown that early elongation relies on a double gradient of JAK-STAT activity controlling apical pulsations ^15^. A fluorescent knock-in of SCAR (SCAR-mNeonGreen) revealed a homogenous expression from one pole to another, suggesting no regulation by the JAK-STAT pathway (Fig S1B) ^26^. Conversely, WRC KD affected neither the gradient of JAK-STAT activity nor apical pulse amplitude (Fig S1A,C). A consequence of pulse gradient is that cells close to the poles are thicker than equatorial cells that are more stretched. In WRC KD, despite the elongation effect, there was still a differential of cell heigh between follicle poles and equator, confirming that pulsatile activity still occurs in gradient, and indicating that cells are still able to stabilize these cell shape changes (Fig S1D). We then wondered whether cell rearrangements could be affected in this context. Although oriented divisions have been observed at stage 5-6, they are not oriented before and there is no more division after stage 6 ^16^. Because the elongation defect in WRC KD appears sooner and carries on afterwards, we reasoned that a potential defect in division orientation cannot be a major contributor to explain elongation failure associated with WRC KD. Finally, we checked oriented cell intercalation. The presence of such events can be estimated between stage 6, during which cells stop dividing, and stage 8, by counting the number of cells in a line going from one pole to the other ^15^. As it is in post-mitotic conditions, an increase in this number necessarily implies cell intercalation. If the number of cells was reported as a function of follicle size, WRC KD almost blocked cell incorporation in the line, with a slope not statistically different from zero (Fig S1E,F). However, if cell number per line was compared with follicle aspect-ratio, their relationship was not significantly different in WRC KD and wild type (WT) situations and both were still different from zero (Fig S1G,H). These data indicate that WRC is required for effective oriented cell intercalation and suggest a direct causal link between intercalation and follicle elongation.

We also looked at follicle cells aspect in epithelium plane at stage 7-8 in wild type and WRC KD and noticed topological differences. Especially, whereas in WT conditions the epithelium shows a regular organization close to the classical honeycomb figure, we detected an accumulation of 4-way vertices in WRC KD (Fig S1I). Such accumulations are typical of a defect in the cell intercalation process, and more precisely of defective new junction formation ^8,10,11^. Since the global pattern of JAK-STAT dependent pulses and cell shape changes seemed unaffected in WRC KD, its impact on cell intercalation was probably due to a more local involvement. To check this assumption, we analyzed mutant clones for null alleles of two WRC subunits, SCAR and Cyfip. These clones clearly revealed an accumulation of 4 way-vertices with three times more of this figure than in WT cells (Fig 1F-H). Even more strikingly, mutant clones showed an accumulation of rosettes, which were almost never observed in the WT situation in the follicular epithelium (Fig 1F,G,I). Thus, altogether, these data show that WRC is required, at tissue scale, for follicle elongation and, at more local scale, for cell intercalation resolution.

### WRC generates a subpopulation of F-actin at tricellular junction

To obtain insight on WRC function in follicle cells, we analyzed its subcellular localization using SCAR-mNeonGreen fluorescent knock-in line. In the most basal plane, SCAR localized as expected in the planar polarized protrusions driving follicle rotation (Fig 2A, Sup movie 1)^25^. Then, on the whole cell height, except for the most apical plane where it was all around the cell, SCAR was found in intense spots corresponding to tricellular junctions (TCJs) (Fig 2A and Sup movie 1). This TCJ localization was observed during all stages of early elongation (Fig 2A and S2A). This enrichment was easily demonstrated by making the intensity ratio between tricellular and bicellular junction and is found both at the levels of adherens junction (E-Cadherin plan) and of more lateral junctions stained with Coracle (Cora) (Fig 2B-D). An identical pattern was observed with a transgene expressing Abi, another WRC subunit, fused to mCherry (Fig 2E). Thus, we concluded that WRC is strongly enriched at TCJs in the follicular epithelium at stages of early elongation. WRC at TCJs colocalized with an intense spot of F-actin (Fig 2E). We developed an automatized detection method of these F-actin spots at TCJs which revealed that they were detectable in their vast majority (Fig 2G and Fig S2B). Importantly, in mutant clones for *SCAR*, these spots completely disappeared (Fig 2F,G). Moreover, this actin enrichment was also lost in mutant clones for *Arp3*, an Arp2/3 complex subunit, indicating that WRC effect was due to its ability to activate this branched actin nucleator (Fig 2G and S2C). Thus, these data establish WRC as a new molecular actor at TCJ where it generates a specific sub-population of branched F-actin.

**Figure 2.**
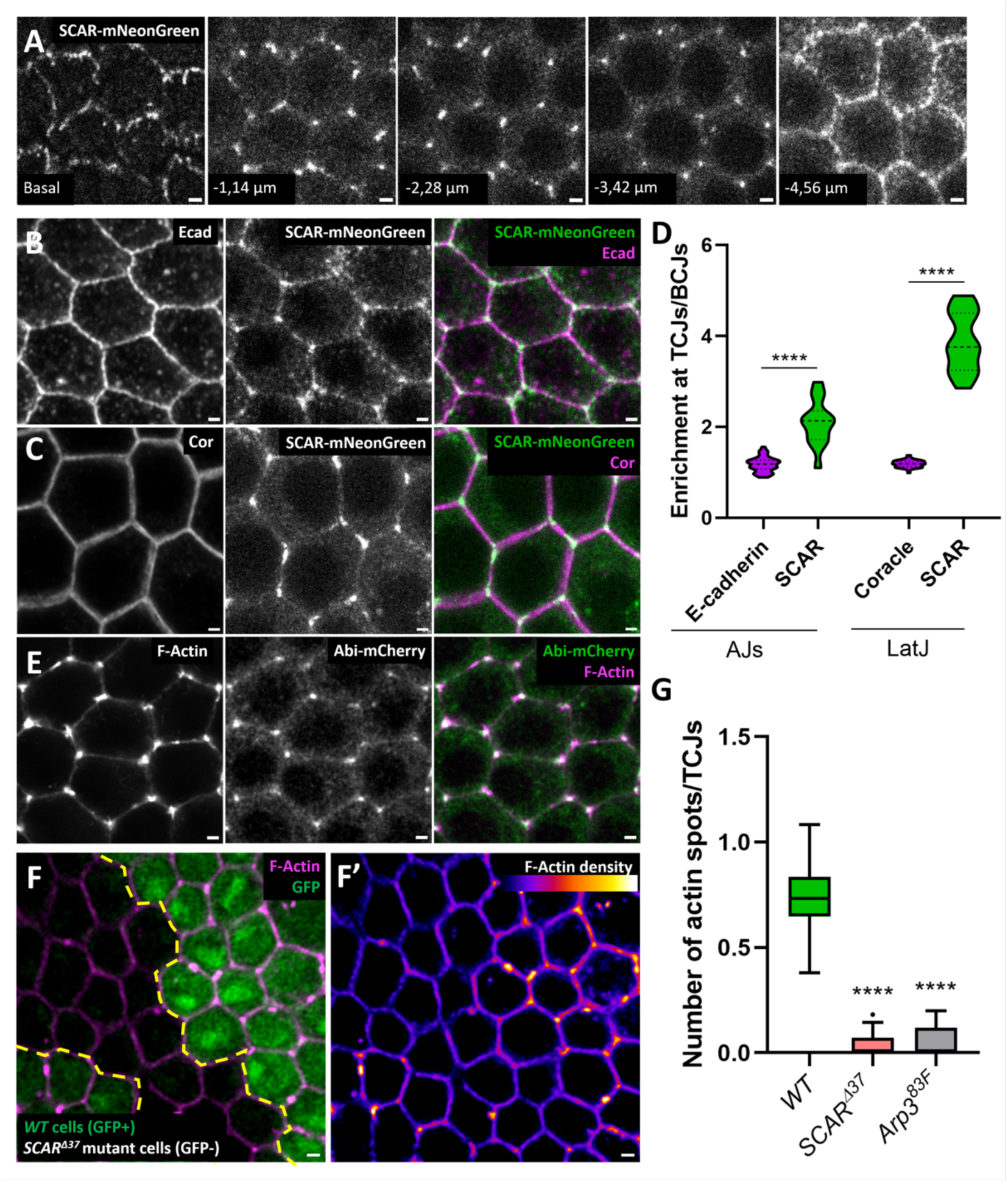
WRC is localized at tricellular junction where it generates a subpopulation of F-actin. A) Snapshots of SCAR-mNeonGreen from a z-stack going from the basal to the apical domain of stage 7 follicle cells. B,C) SCAR-mNeonGreen is observed at tricellular junctions both in the plan of adherens junction, marked with Ecad (B) and of septate junctions marked with Cora (C). D) SCAR-mNeonGreen intensity quantification of tricellular versus bicellular junctions at the level of adherens (AJs) and lateral junctions (LatJ). E) Abi-mCherry is detected at TCJ where it co-localizes with an intense spot of F-actin. F) mutant clone for *SCAR^◿^*^37^ marked by the absence of GFP and stained for F-actin. F-actin intensity is color-coded on F’. G) Quantification of the F-actin spot number per TCJ in the indicated genotypes. (Scale bars: 1 μm)(**** P < 0.0001)

### WRC induces dynamic lamellipodia invading bicellular junctions

We decided to better characterize this population of TCJ F-actin. High resolution image revealed that it was not always an isotropic spot, but rather that intense F-actin staining frequently extended on a bicellular junction (Fig 3A). This actin population, when occurred, was usually on only one of the three bicellular junctions joined at a TCJ and owned a maximal length of about 1.5 µm, though longer ones could be unfrequently observed (Fig 3B). Given that WRC usually induces protrusion, we wondered whether it could be also the case at TCJ. Clones of cells expressing LifeAct-RFP were induced and TCJ at clone boundaries were analyzed. Strikingly, cells expressing LifeAct-RFP sent actin rich protrusions from TCJs into the bicellular junctions between pair of cells that did not express LifeAct-RFP (Fig 3C). On the other hand, we did not detect actin enrichment on the bicellular junction in between two cells expressing LifeAct-RFP but in front of a cell that did not express it. These results indicate unambiguously that TCJ F-actin corresponds to a bona fide protrusive activity.

**Figure 3.**
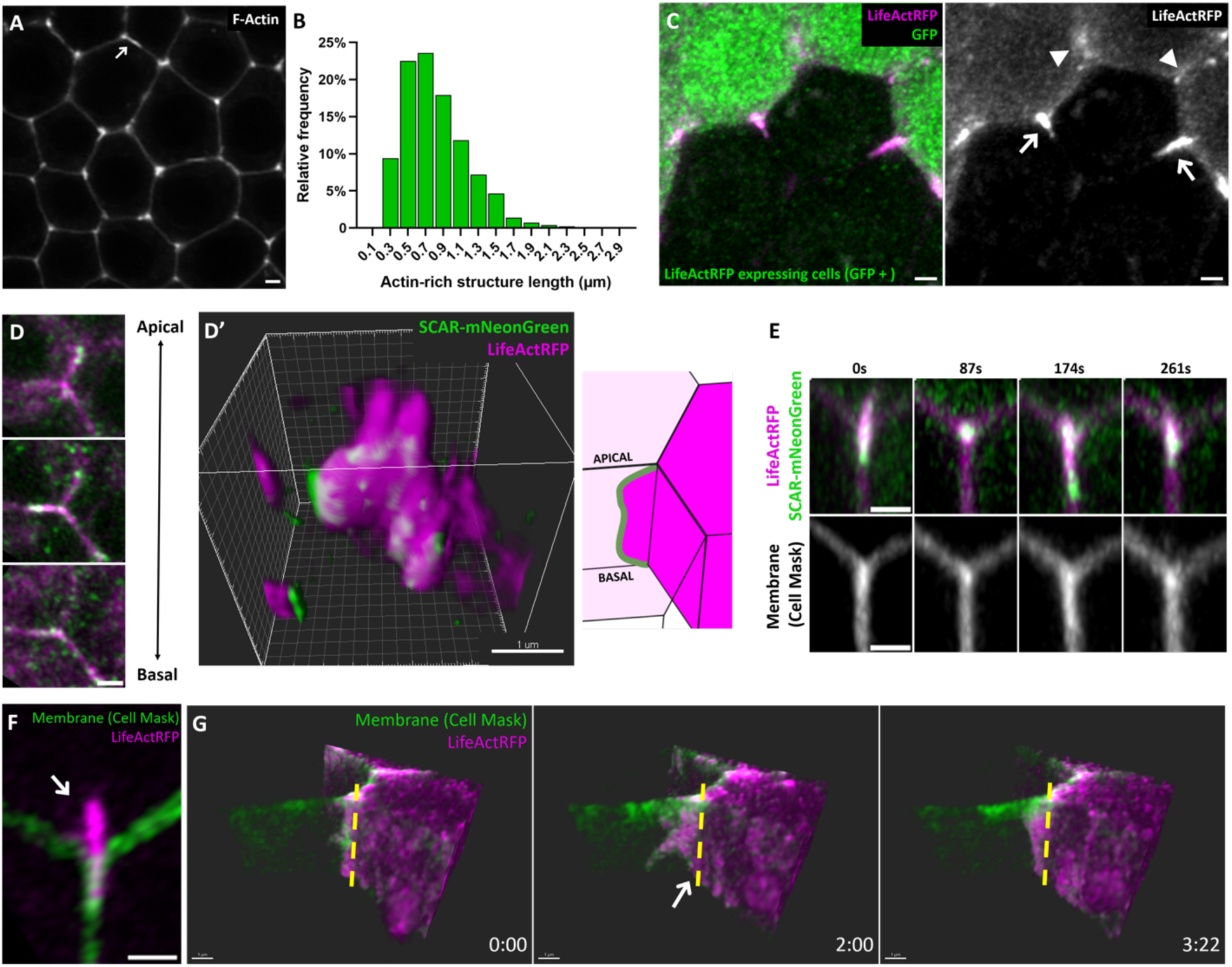
F-actin dynamics at tricellular junction. A) High resolution image of F-actin in a stage 7 follicular epithelium. Arrow shows a bright signal extending on a bicellular junction. B) Distribution of F-Actin intense signal at TCJ according to its length. C) Fixed image of a mosaic follicle in which only GFP-positive cells express LifeAct-RFP. Arrows indicate protrusive activity from an expressing cell in between two non-expressing cells and arrowheads the absence of F-actin rich structure in front of non-expressing cells. D) Snapshots of SCARmNeonGreen and LifeAct-RFP from a z-stack going from the basal to the apical of follicle cells and stained for F-actin. D’) 3D reconstruction of the protrusion from D. E) Snapshots from a movie at a TCJ in a SCARmNeonGreen follicle expressing LifeAct-RFP and stained with CellMask. F) Snapshot from a protrusive activity at TCJ forming a whip-like structure in the cytoplasm. H) Snapshots from a 3D reconstruction of a movie on a follicle with clonal expression of LifeactRFP and stained with CellMask. Dashed line indicates the position of the tricellular junction (Scale bars: 1 μm).

WRC is known to mainly induce lamellipodia rather than other kinds of protrusion such as filopodia. After F-actin staining on SCAR-mNeonGreen follicles, we performed 3D reconstruction of the protrusion induced from a TCJ. This reconstruction clearly illustrated that protrusion could spread on almost the whole cell heigh of the cells from apical to basal, thus corresponding to lamellipodia (Fig 3D and supplemental movie 2). As expected, SCAR localized at the front of this lamellipodia where it induced actin polymerization (Fig 3D). We also performed movies using LifeAct-RFP and SCAR-mNeonGreen and a fluorescent dye for plasma membrane to follow the dynamics of this protrusive activity (Fig 3E and supplemental movie 3). Protrusion appeared and disappeared in about a minute and two successive protrusions could be observed in about 3 minutes. SCAR, moving from the TCJ, was observed at the front of the protrusion, where WRC is expected to induce polymerization. Moreover, the membrane marker was brighter at protrusion position, confirming the double amount of membrane due to the protrusive activity between adjacent cells (Fig 3E). Notably, actin polymerization was sometimes faster than protrusion extension, maybe because of the resisting force of the bicellular junction, and led to the formation of comet or whip-like structures extending in cell cytoplasm (Fig 3F and supplemental movie 4). Altogether, these data indicate that the WRC-dependent F-actin subpopulation at TCJ is protrusive, highly dynamic and forms lamellipodia in between adjacent cells. These properties can be summarized by the 4D visualization of such a lamellipodia on live samples with clonal expression of LifeAct-RFP (Fig 3G and supplemental movie 5).

### WRC promotes intercalation resolution from cells at extremities of the new junction

We tried to connect WRC molecular activity, protrusive actin polymerization, with its impact at supracellular level, which is to promote cell intercalation resolution. To directly probe a potential link between WRC activity and new junction formation, we analyzed actin intensity on short junctions. We noticed that short junctions (less than 1.5 µm) could be fully covered by intense F-actin associated with SCAR (Fig 4A). However, there was no bias in the probability to observe a protrusion on a junction depending on its length (Fig 4B). Nonetheless, if considering junctions with lamellipodia, the probability to have a junction fully covered by a protrusion was strong for junction inferior to 1 µm and logically dropped very fast for longer junctions, as protrusion length rarely exceeds 1.5 µm (Fig 4C). Thus, if WRC-dependent protrusive activity is associated with the appearance of new junctions, its impact is likely limited to the very first step of elongation.

**Figure 4.**
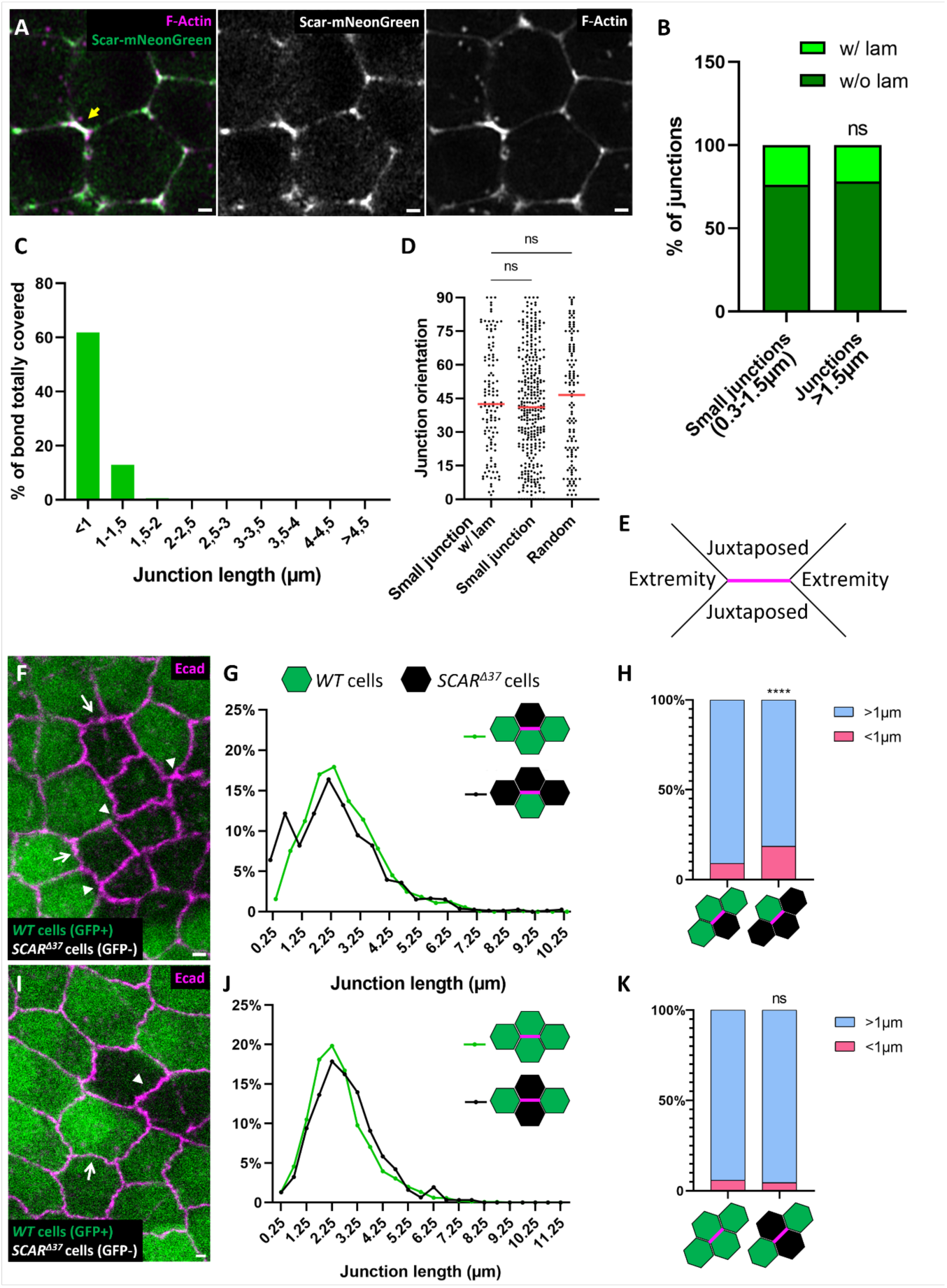
WRC promotes intercalation resolution and junction lengthening from extremity cells. A) Image of a stage 7 SCAR-mNeonGreen follicle stained for F-actin. Arrow shows a small junction strongly enriched for both SCAR and F-actin. B) Proportion of bicellular junction with (w/) our without (w/o) lamellipodia. Two classes were analyzed: short junctions (between 0.3 and 1.5 μm) and longer ones (>1.5 μm). C) Among junctions associated with a lamellipodia, proportion of them fully covered by a protrusion depending on their length. D) Angular distribution of total short junctions, short junctions enriched in F-actin and compared with a random distribution. E) Scheme illustrating the definition of “juxtaposed cells” and “extremity cells” of a given junction. F,I) Follicles with mutant clones for *SCAR^◿^*^37^ marked by the absence of GFP and stained for E-cad. Arrows show junctions for which F) extremity cells or I) juxtaposed cells are wildtype and arrowheads junction for which F) extremity cells or I) juxtaposed cells are mutants for *SCAR*. G,J) Distribution of junction length according to the genotype of G) extremity cells and J) juxtaposed cells (WT or mutant for *SCAR*). H,K) Proportion of short junctions (<1m) depending on the position of *SCAR* mutant cells (Scale bars: 1μm )(**** P < 0.0001).

As the cell intercalations are globally polarized in the follicular epithelium, we analyzed the angular distribution of short junctions to detect an eventual preferential orientation. We analyzed all the short junctions and the ones enriched in F-actin and compared their angular distribution with a randomized distribution (Fig 4D). No preferential orientation was observed in any case, although we could not exclude that it may occur in regions of the follicle that cannot be easily analyzed such as follicle poles. This observation suggests that WRC protrusive activity promotes tissue elongation by permitting cell rearrangements and providing enough tissue plasticity rather than by strictly controlling the orientation of these rearrangements.

Accumulation of 4-way vertices and rosettes, that we observed in WRC mutants, is usually associated with a defect in lengthening of new junctions. Such defect is observable on fixed samples by a change in the distribution of junction length with an accumulation of short junctions^8^. We therefore analyzed this distribution in mutant clones for SCAR. Since WRC owns a protrusive activity that spreads in the bicellular junction of two juxtaposed cells, we reasoned that it might be specifically required in the cells at new junction extremities (extremity cells). We therefore refined our analysis on clone borders and reported junction distribution depending on the genotype of cells at junction extremities (Fig 4F,G,H). If extremity cells were WT, the distribution was as in control conditions. Strikingly, if extremity cells were both mutants, there was a significant alteration of junction distribution with an accumulation of short junctions, inferior or egal to 1 µm. We also assessed the impact of juxtaposed cells genotype in conditions where extremity cells were wild type (Fig 4 I,J,K). No differences in junction length distribution between wild type and SCAR mutant cells were observed. Altogether, these data demonstrate that WRC promotes short junction lengthening specifically in cells at junction extremities, in agreement with its protrusive activity.

### WRC can be recruited by Sidekick

Our data strongly suggested that WRC at TCJ was critical for epithelium dynamics but how WRC was recruited at this very specific location was unknown. It has been proposed that WRC can interact at cell cortex with transmembrane proteins containing a specific motif called WIRS for WRC interacting receptor sequence ^27^. We therefore looked for transmembrane proteins with a conserved WIRS motif from fly to human and expressed in follicle cells. Four proteins correspond to these criteria and, strikingly, Sidekick was one of them. Endogenous GFP-tagged Sdk strongly colocalized with both WRC (Abi-mCherry) and F-actin at TCJ in follicle cells (Fig 5A). Of notice, in follicle cells, Sdk was found both at the levels of subapical adherens junction and lateral junction (Fig S3A), probably reflecting the absence of mature septate tricellular junctions at these stages ^28^. We therefore tested the potential interaction between Sdk and WRC by performing coimmunoprecipitation (Fig 5D). WRC subunit HSPC300 fused to GFP was co-expressed in follicle cells in presence of a HA-tagged Sdk transgene wildtype or deleted of its WIRS motif (Sdk^ΔBD^). Sdk-HA but not Sdk^ΔBD^-HA was coimmunoprecipitated with HSPC300-GFP, showing a biochemical interaction depending on its WIRS motif. To confirm this interaction, we also used the GRAB-FP system ^29^. Endogenous GFP-tagged Sdk was delocalized to lateral bicellular junctions using a GFP nanobody fused to Nrv1 (and to BFP for its visualization). In such context, but not in absence of GFP-Sdk, Abi-mCherry was also found on the lateral bicellular domains of the cells indicating that it was trapped by Sdk (Fig 5B,C). Sdk and Abi were also found together with GRAB-FP in big intracellular aggregates (Fig 5B). This experiment confirmed the interaction between the WRC and Sdk, which has also been recently described using different approaches in vitro an in the fly retina (Malin et al., 2022). These data support the idea that Sdk can recruit WRC where it induces actin polymerization. In such a model, Sdk should be found at the tip of WRC-dependent protrusions. We performed movies of GFP-Sdk follicles that also expressed LifeAct-RFP and in which membranes were stained (Fig 5E and supplemental movie 6). Strikingly, when a protrusion appeared, Sdk moved from the TCJ with the protrusion front. Thus, Sdk can recruit WRC and its dynamics fulfills the classical scheme for an anchoring membrane partner of the WRC, directing protrusive activity.

**Figure 5.**
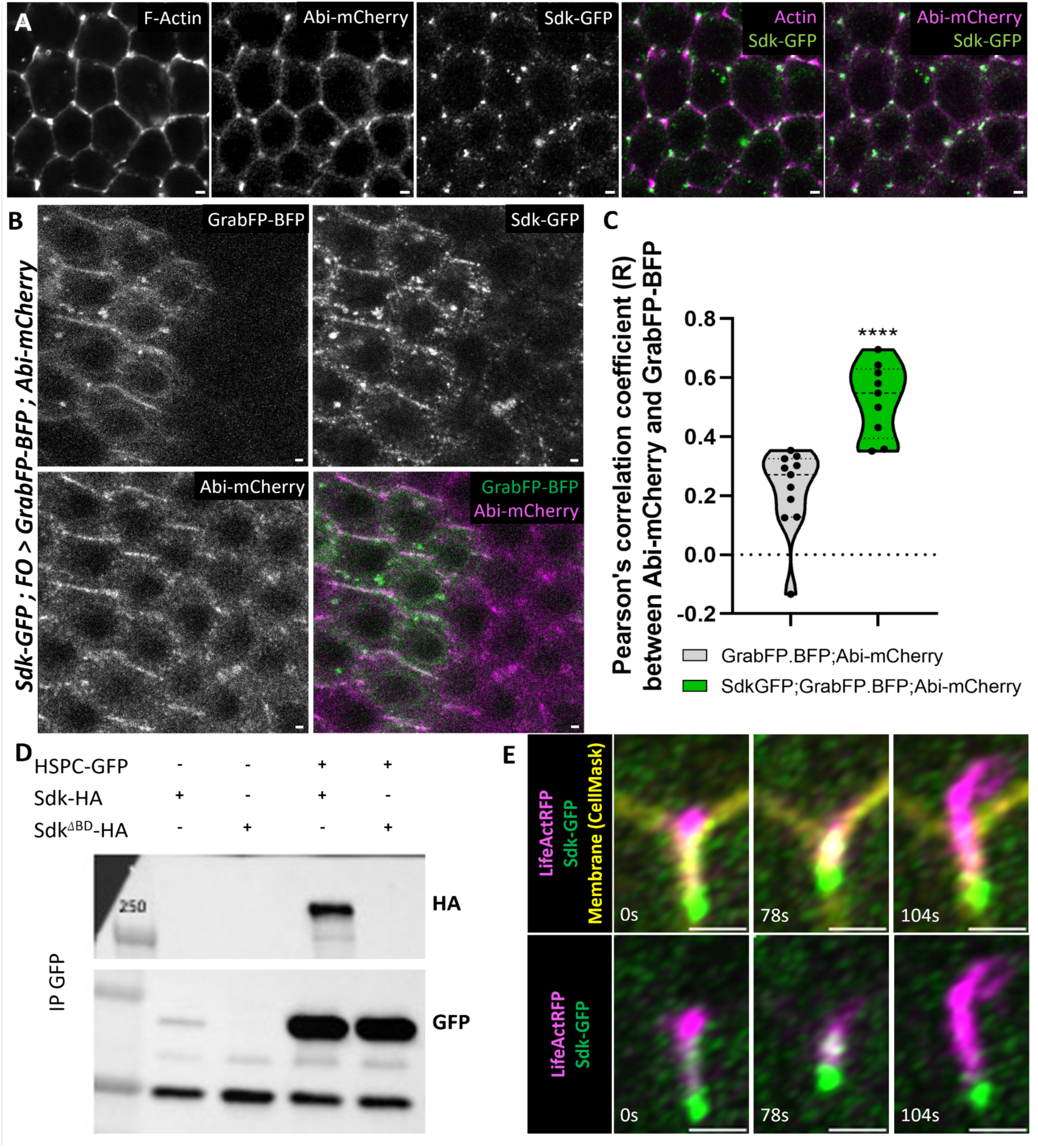
Sdk behaves as a membrane anchor for the WRC. A) stage 7 follicle expressing Abi-mCherry and endogenous Sdk-GFP, and stained for F-actin B) Images from a follicle with mosaic expression (obtained by flip-out experiment (FO)) of lateral GrabFP fused to BFP, in presence of Abi-mCherry and endogenous Sdk-GFP. C) Quantification of colocalization between GrabFP and Abi-mCherry in presence or absence of Sdk-GFP using Pearson’s coefficient. Measures were made in ROIs corresponding to high level of GrabFP areas (i.e. lateral junctions and aggregates) based on intensity thresholding. D) Western blots against HA (top) and GFP (bottom) after co-immunoprecipitation against GFP on ovaries expressing the indicated transgenes in the follicle cells. E) Snapshots of a movie of Sdk-GFP follicle expressing LifeAct-RFP and stained with a membrane dye (Scale bars: 1μm)(**** P < 0.0001).

### WRC is redundantly recruited by Lar and Sdk at tricellular junctions

Our data fits with a model in which Sdk recruits WRC and, when the latter is activated, they generate together a protrusion from the tricellular junction that extends into a bicellular junction. Moreover, intercalation defects due to the loss of WRC in follicle cells are highly reminiscent of the ones observed in *sdk* mutant in other tissues, in accordance with the hypothesis that they cooperate to resolve cell intercalation ^10–12^. However, in *sdk* null mutant follicles, WRC was still present at TCJ and F-actin spots and protrusions were still observed (Fig S6A and Fig 6C). Moreover, these follicles elongated properly, and we did not detect any significant topological defect (Fig 6K, 6G-H and Fig S6D). We therefore hypothesized that Sdk could be redundant with another protein in its ability to recruit WRC at the TCJ.

**Figure 6.**
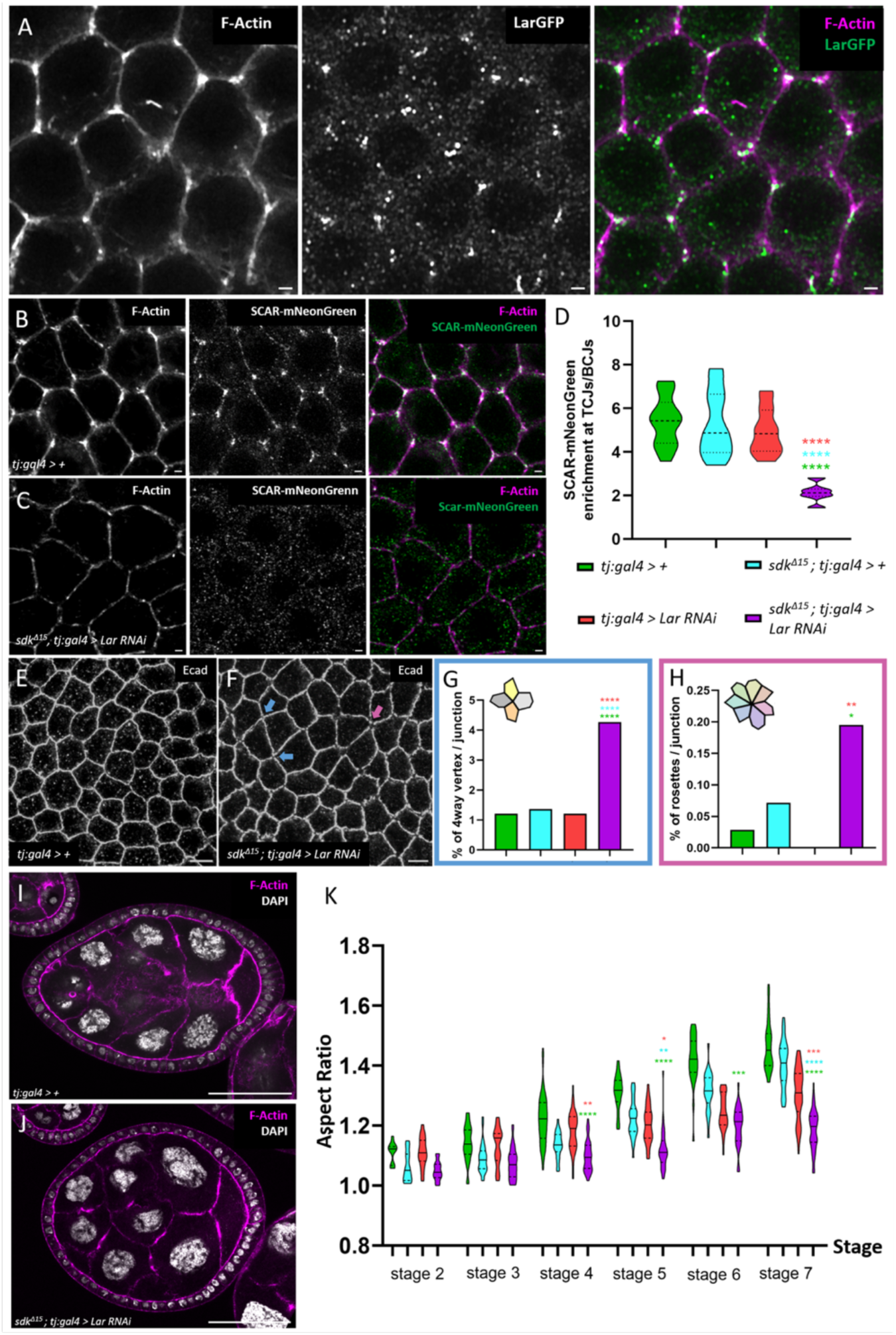
Redundancy between Lar and Sdk for WRC recruitment, cell intercalation resolution and tissue elongation. A) Colocalization between F-actin spots and endogenous Lar tagged with 3xGFP at TCJs. B-C) Scar-mNeonGreen and F-actin in B) control follicle and C) follicle mutant for *sdk* and with RNAi against *Lar*. D) Quantification of the presence of Scar-mNeonGreen at TCJs in the indicated genotypes. E-F) Immunostaining against E-Cad in a E) control follicle and F) follicle mutant for *sdk* and with RNAi against *Lar*. Blue arrows show 4-way vertices, pink arrow shows rosette. G-H) Quantification of 4-way vertices (G) and rosettes (H) frequencies in the indicated genotypes. I,J) Sagittal view of a I) control follicle and J) follicle mutant for *sdk* and with RNAi against *Lar*, stained with F-actin and DAPI. K) Aspect-ratio quantification depending on stage and of the indicated genotypes. (Scale bars: 50μm of whole follicle images, 1μm on follicle cell images) (* P < 0.05, ** P < 0.01, ***P < 0.001, **** P < 0.0001).

Lar is a receptor tyrosine phosphatase that can interact with WRC and induce lamellipodia on the basal surface of follicle cells ^22,30^. Observation of endogenous Lar tagged with 3xGFP clearly showed that Lar was enriched at tricellular junctions where it colocalized with F-actin (Fig 6A) ^31^. However, although Lar is involved in follicle rotation and the subsequent elongation, mutant follicles for *Lar* elongated almost normally during early stages (Fig 6K and S6G). Moreover, F-actin at tricellular junctions and 4-way vertex number were similar to WT (Fig S6B,C,E and Fig 6G). We therefore tested a potential redundancy between Lar and Sdk.

We first tried to obtain double mutant flies for *sdk* and *Lar*, as both mutations are individually homozygous viable. However, only very few adult flies of this genotype were recovered, suggesting a synthetic sublethality between those two genes. Moreover, these flies died quickly after birth precluding proper analysis of their ovarian follicles. We therefore induced *Lar* KD specifically in follicle cells by RNAi in a *sdk* mutant background in flies with SCAR-mNeonGreen knock-in. In such context, SCAR was no longer enriched at TCJs, whereas its localization was not affected in *sdk* mutant or in *Lar* KD (Fig 6B-D and Fig S6A-B). Accordingly, TCJ F-actin subpopulation was also fully compromised (Fig 6B,C and Fig S6A-C). Thus, Sdk and Lar proteins are redundant for the recruitment of WRC at TCJ. Hence, this genetic combination presents a unique opportunity to confirm the role of this specific F-actin population in cell and tissue dynamics. Concomitant loss of *sdk* and *Lar* affected cell topology with 4-way vertices and rosettes accumulation and tissue shape with a strongly reduced follicle elongation during early stages (Fig 6E-6K). Thus, blocking WRC recruitment at TCJ reproduces its loss of function at molecular, cellular and tissular scales, confirming the causal links between protrusive activity at TCJ, cell intercalation resolution and tissue elongation.

## Discussion

This work shows that a specific subpopulation of F-actin emanates from tricellular junctions and that it is critical for epithelium dynamics and morphogenesis. While myosin II impact on cell intercalation is widely recognized, a role for F-actin polymerization was more elusive. F-actin spots at TCJs have been detected in other epithelia, as in fly embryo ectoderm, where their presence is dependent on Sdk ^12,32^. Perhaps more significantly, a very similar pattern was, for instance, observed in the vertebrate neural plate ^33,34^. Thus, this actin population is likely relevant for the development of many animal epithelia, revealing a new key player at TCJ.

Although it has not been connected to TCJs, lateral protrusive activity has been noticed during the development of different epithelial tissues. So far, such protrusions have been proposed to promote junction shrinkage rather than intercalation resolution and the subsequent junction lengthening. For instance, during germband extension of fly embryo, lateral protrusions containing Rac and PIP3, two WRC activators, precede rosette formation ^35^. In *C. elegans*, lateral protrusions, redundantly generated by WRC and WASP (another Arp2/3 activator), are required to initiate intercalations of the dorsal epidermal cells ^36^. Lateral protrusions have been also described in the mouse neural plate, fitting with the previously mentioned actin spots at TCJ in this tissue, although neither the origin nor the precise function of these protrusions have been analyzed ^37^. Our data reveals that, in follicle cells, WRC activity is also required for intercalation resolution by promoting the lengthening of short junctions. Importantly, blocking WRC recruitment at TCJ reproduces these defects, showing that it is specifically this actin subpopulation that is mandatory.

How this actin population with a protrusive activity promotes lengthening of short junctions in the follicle cells will need further investigation in the future. In fly male genitalia, Sdk promotes intercalation resolution and junction lengthening by favoring cortical Myosin II enrichment in extremity cells compared to the new junction between juxtaposed cells ^10^. Quite similarly, both WRC and Sdk have been involved in junction lengthening in the fly retina by limiting MyoII recruitment on this junction ^21,38^. Thus, in each case, TCJ dynamics allows the lengthening of short junctions by creating a bias in cortical tension. Nonetheless, in the fly retina, the possibility that WRC is required in a non cell-autonomous manner, i.e. from the equivalent of the extremity cells, has not been explored. WRC is typically known to induce protrusive F-actin population, as, for instance, in lamellipodia of migrating cells. Moreover, our data showing its presence at protrusion tip, recruited by transmembrane proteins as Lar and Sdk also correspond to its classical mode of action (Chen et al., 2014). Our genetic analysis of mosaic tissues showing that WRC is required in the extremity cells is coherent with its protrusive activity. Of notice, average protrusion length is in the same range as that of accumulating short junctions if protrusive activity is blocked. In other words, this protrusive activity likely explains the specific behavior of short junctions described in different contexts, as during cell intercalation resolution or in very small cells as in the fly retina ^8,10,11,38^. Since in different tissues WRC subunits or *sdk* loss of function induces similar defects than in follicle cells and that Sdk can recruit WRC, we propose that WRC protrusive activity from TCJ promoting junction lengthening in a non cell-autonomous manner is a common mechanism of epithelium morphogenesis. Moreover, the involved motif in Sdk protein is found in almost all the animal kingdom, suggesting that this interaction reveals a conserved mechanism ^39^. Finally, a recent publication has also reported a molecular interaction between WRC and Sdk in the fly retina using different approaches than the ones described here, giving together very robust evidence for this interaction ^21^. However, only a weak diminution of SCAR junction enrichment in *sdk* mutant retina was described, leading authors to propose a potential redundancy for WRC recruitment. In the follicle cells, *sdk* mutation induces no defect on its own, though Sdk delocalization shows that it can recruit WRC. Nonetheless, concomitant loss of function of Sdk and Lar fully abolished WRC recruitment at TCJ. Importantly, it also induces cell intercalation and tissue elongation defects confirming that actin dynamics at TCJ is essential for tissue dynamics at different scales.

Thus, our work identifies Lar as a functional component of TCJ involved in cell rearrangements and demonstrates a redundancy between Lar and Sdk. Moreover, the synthetic sublethality between null mutants for *sdk* and *Lar* suggests that this redundancy is relevant for other tissues, explaining, by the way, the limited impact of single mutant. We detected no bias in the orientation of F-actin enriched short junctions, suggesting no spatial control while oriented cell intercalations in fly follicle cells occur. If so, protrusive activity at TCJ would be permissive for intercalations, providing fluidity to the tissue. However, Sdk has been proposed to be part of a mechanosensitive mechanism while Lar function has been associated in epithelial cells with planar cell polarity ^12,30^. Thus, WRC activity might be part of an instructive mechanism relying on different spatial cues, i.e. biochemical and/or mechanical. Further investigations will be therefore required to determine whether their molecular redundancy reflects distinct mechanisms for the proper spatiotemporal control of WRC activity, allowing robust morphogenesis when they work in combination.

Nonetheless, this work highlights the relevance of actin dynamics at tricellular junction for cell intercalations, and more specifically for their resolution and the initiation of a new junction, and provides important data on how this specific protrusive population is generated through the recruitment of WRC.

## Methods

### Drosophila Genetics

Fly stocks with their origin and reference are described in Supplementary Table 1. Full genotypes and temperatures for each experiment are provided in Supplementary Table 2. Crosses were raised at 25°C and experimental females were aged 3-5 days with males and put on yeast 1 day before dissection, unless specified in Supplementary Table 2. For clonal induction and flip-out experiments, two heat-shocks over 2 days were performed at 37°C during 1h on adult flies.

### Microscopy and live imaging

Ovaries dissection and immunostaining were performed as previously described (Vachias et al., 2014) in supplemented Schneider. Exceptions are for F-Actin staining which was performed after 15min PFA 16% fixation and permeabilization with PBS + 0.02% Triton. Images were taken using Zeiss LSM800 Airyscan or Zeiss LSM980 Airyscan. Stage determination was performed using unbiased criteria of nurse nuclei diameter (Dai et al., 2017), with adjustments for stage 6/7 based on cell divisions. Primary antibodies used were against DE-Cad (DHSB AB_528120 ; 1/200), Cora (DHSB AB_1161642 ; 1/200), Lar (DHSB AB_528202 ; 1/200) and secondary antibodies were Donkey anti-rat Cy3 (Jackson 712-165-150 ; 1/500) Donkey anti-rat AlexaFluor488 (Invitrogen A21208 ; 1/500) and Donkey anti-mouse Cy3 (Jackson 715-165-150). F-Actin staining were performed with phalloidin-Atto 488 (Sigma 49409; 1/500), phalloidin-Atto 550 (Sigma 19083) or phalloidin-Atto 633 (Sigma 68825).

For live imaging, individual ovarioles were transferred into a 30μL micro-well (Ibidi BioValey). Samples were cultured less than 2hr before imaging on Zeiss LSM980 with Fast Airyscan module (excepted for Fig S1.C where the imaging was performed on Zeiss Spinning Disk). When used, Deep Red Cell Mask (ThermoFisher C10046) was added in culture medium at a final dilution of 1/5000.

### Co-immunoprecipitation

Co-immunoprecipation was performed on ovary extract from 10 females of each genotype with 100μl of lysis buffer composed of 10mM Tris ph8, 500mM NaCl, 1% NP40, 1mM ortho-vanadate, complete EDTA free antiprotease. Anti-GFP antibodies solution (1:1 anti-GFP 19C8 and 19F7 from Heintz Lab, Rockefeller University) were added at final concentration of 1μg/μL to the extract for 2 hours at 4°C. Protein L Magnetic Beads (Pierce 88850) were equilibrated with 10 volumes of 10mM Tris ph8, 500mM NaCl, 1% NP40, 1mM ortho-vanadate, 1% BSA. Then, 100μl of this solution was added in new tubes and the supernatant eliminated before adding the ovary extract during 30 minutes at 4°C. Beads with immunocomplexes were washed with 10mM Tris phH8, 500mM NaCl, 1%NP40. Western blot was done using standard procedures using anti-HA antibody (Roche 11867423001) and anti-GFP antibody (JL-8, Clontech).

### Image analysis

Aspect Ratio and Elongation coefficient were calculated based on measure of the length of the long and short axis of each follicle as previously described using Fiji software (Alégot et al., 2018).

4-way number was automatically determined after cell segmentation based on Ecad staining using TissueAnalyser software with a threshold of inclusion of less than 0,3μm junction length (Aigouy et al., 2016). Rosette count was performed manually based on Ecad staining.

Enrichment at tJs/bJs was performed as described in Letizia et al., 2019 after cell segmentation based on Ecad or Cora stainings. AJs section correspond to a Z-max projection of 1μm to 2μm around maximal Ecad plan. LatJs section correspond to a Z-max projection of 1μm in Cora plan.

Actin-rich population detection and parameters measurements were done using homemade FIJI macro based on F-Actin intensity thresholding on segmented images, allowing to determine their association with TCJ and their length. For figure 4, a lamellipodia was defined as an actin spot with an aspect ratio superior or equal to 1.5.

3D reconstructions of lamellipodia were done using IMARIS software.

### Statistical analysis

For all experiments, sample size is indicated in Supplementary Table 3. Results were obtained for at least two independent experiments. For each experiment, at least five females were dissected. The experiments were not randomized or blinded. Statistical analysis and graphs were made using GraphPad Prism. Briefly, after verification of normal distribution using Shapiro test, statistical significance was assessed by ANOVA with Tukey’s post-hoc test for multiple comparisons and by Student’s t test for comparison between two groups. For elongation coefficient and linear regression of cell numbers in sagittal views, statistical significance was tested by ANCOVA. Differences in junction length distribution or 4-way vertices and rosettes proportion were tested by Chi-square. For all figures, p*<0.05, **<0.01, ***<0.001, ****<0.0001.

## Acknowledgments

We thank Greg Bachaw, Jenny Gallop, Angela Giagrande, Sally Horne-badovinac and Jessica Treisman for sharing fly stocks. Stocks obtained from the Bloomington Drosophila Stock Center (NIH P40OD018537) were used in this study. We thank Yohanns Bellaïche and Jesus Lopes-Gay for exchanges. LC was supported by Fondation pour la Recherche Médicale (FRM) and Association pour la Recherche contre le Cancer (ARC) grants. HA was supported by the Region Auvergne Rhône Alpes. VM obtained financial support from Ligue Contre le Cancer. This research was also financed by the French government IDEX-ISITE initiative 16-IDEX-0001 (CAP 20-25). We also thank the CLIC facility (Clermont Imagerie Confocale) for technical support, and team members for comments on the manuscript.

---

**Figure S1.**
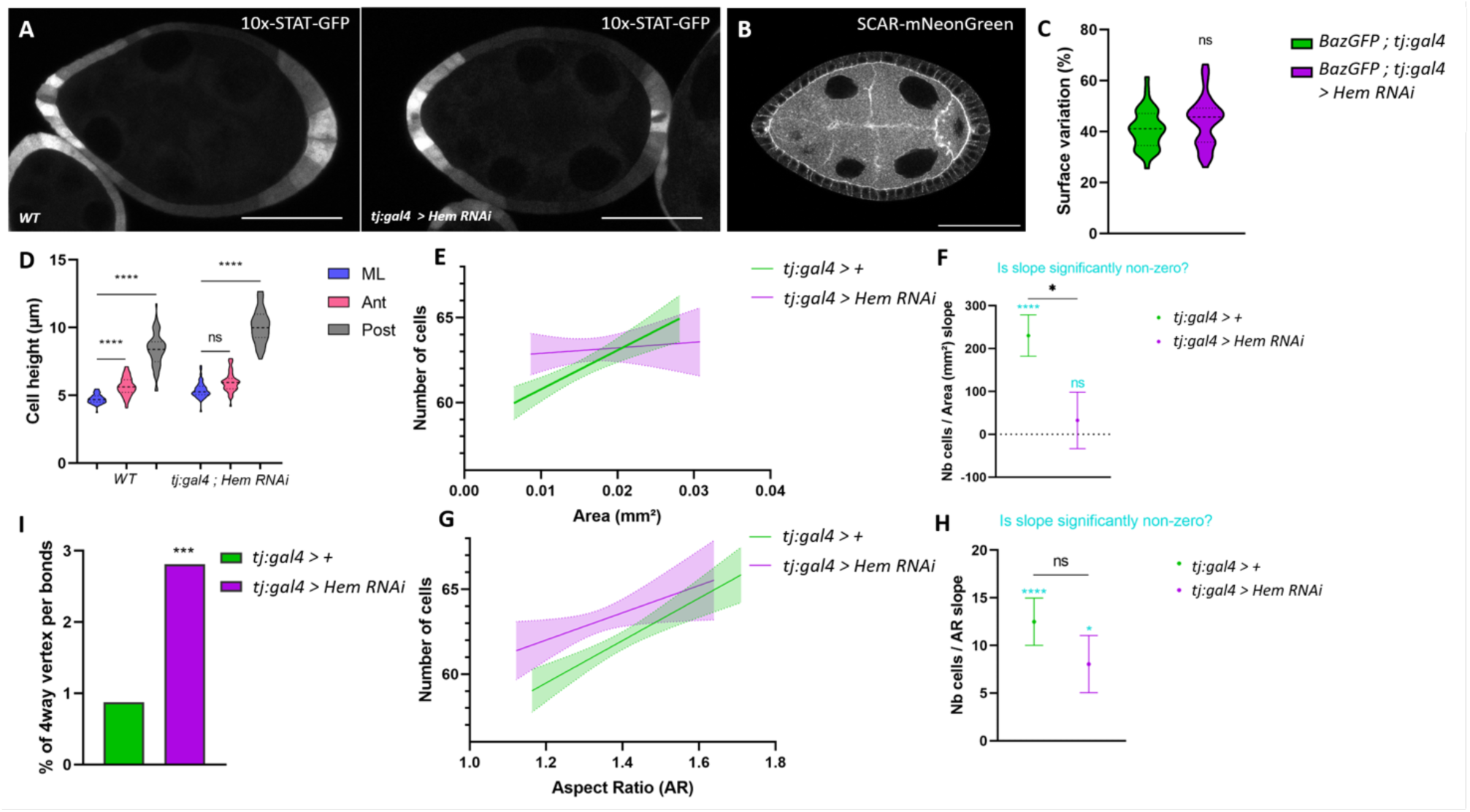
A) Reporter of Jak-STAT activity (10x-STAT-GFP) in control and *Hem* KD conditions. B) Sagittal view of stage 7 follicle from the *SCAR-mNeonGreen* knock-in line. C) Analysis of apical pulse amplitude (surface variation) in control and *Hem* KD conditions. D) Measure on stage 7-8 follicles of cell heigh in the mediolateral region (ML) or close to the poles (Ant and Post). E,G) Number of cell on a sagittal plan from one pole to the other of postmitotic 6-8 stage follicles as a function of E) follicle size (area of the plan used for cell counting) or G) aspect ratio (AR) in control and Hem KD conditions. F,H) Representation of the slopes from E,G and statistical comparison to a null slope. I) quantification of the number of 4 way-vertices per junction in control and *Hem* KD conditions on stage 7-8 follicles. (Scale bars: 50μm)( ** P < 0.01, ***P < 0.001, **** P < 0.0001).

**Figure S2.**
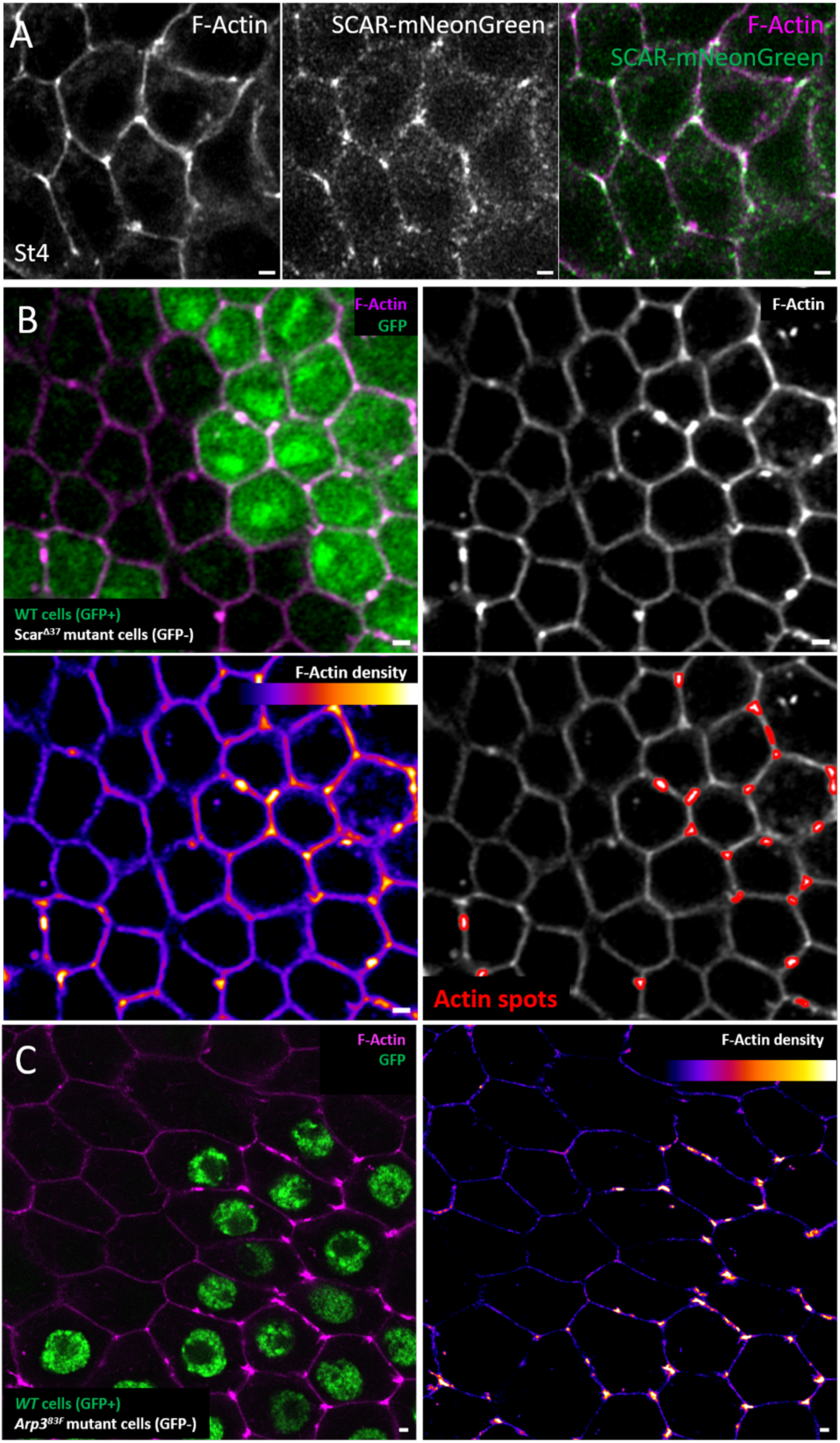
A) Stage 4 follicle from SCARmNeonGreen knock-in line and stained for F-actin. B) Same mutant clone for *SCAR^◿^*^37^ marked by the absence of GFP and stained for F-actin as shown on fig 2F. Last panel illustrates the automatic detection of F-actin spots. C) Image of mutant clone for *Arp3^83F^*marked by the absence of GFP stained for F-Actin. (Scale bars: 1 μm).

**Figure S5.**
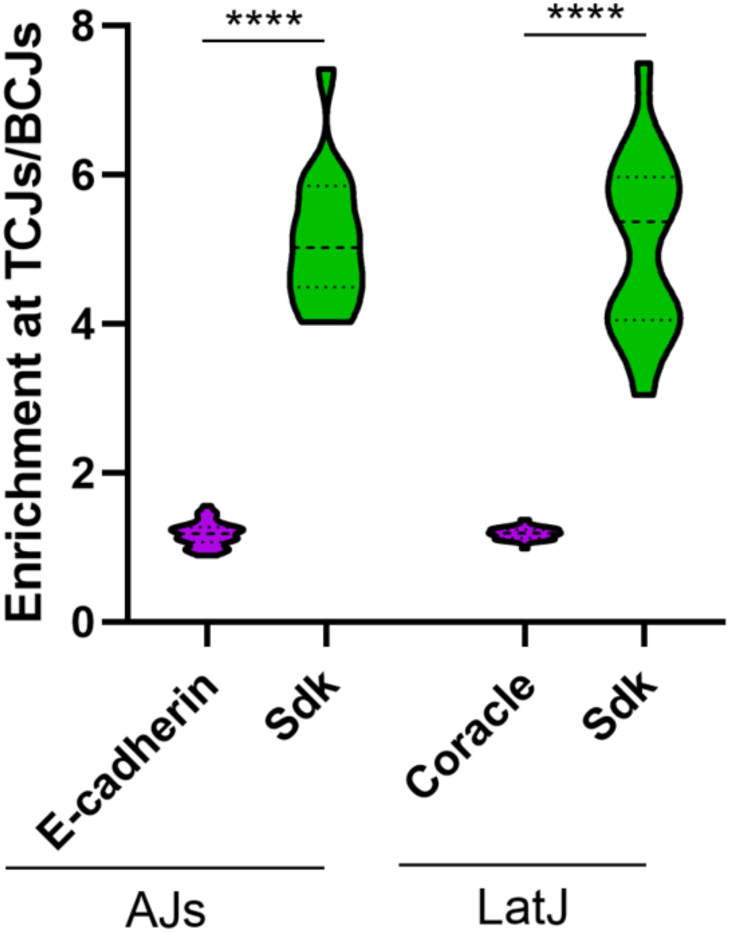
Quantification of tricellular versus bicellular enrichment of Sdk at the level of adherens junctions (compared to E-cad) and lateral junctions (compared to Cora) (**** P < 0.0001).

**Figure S6.**
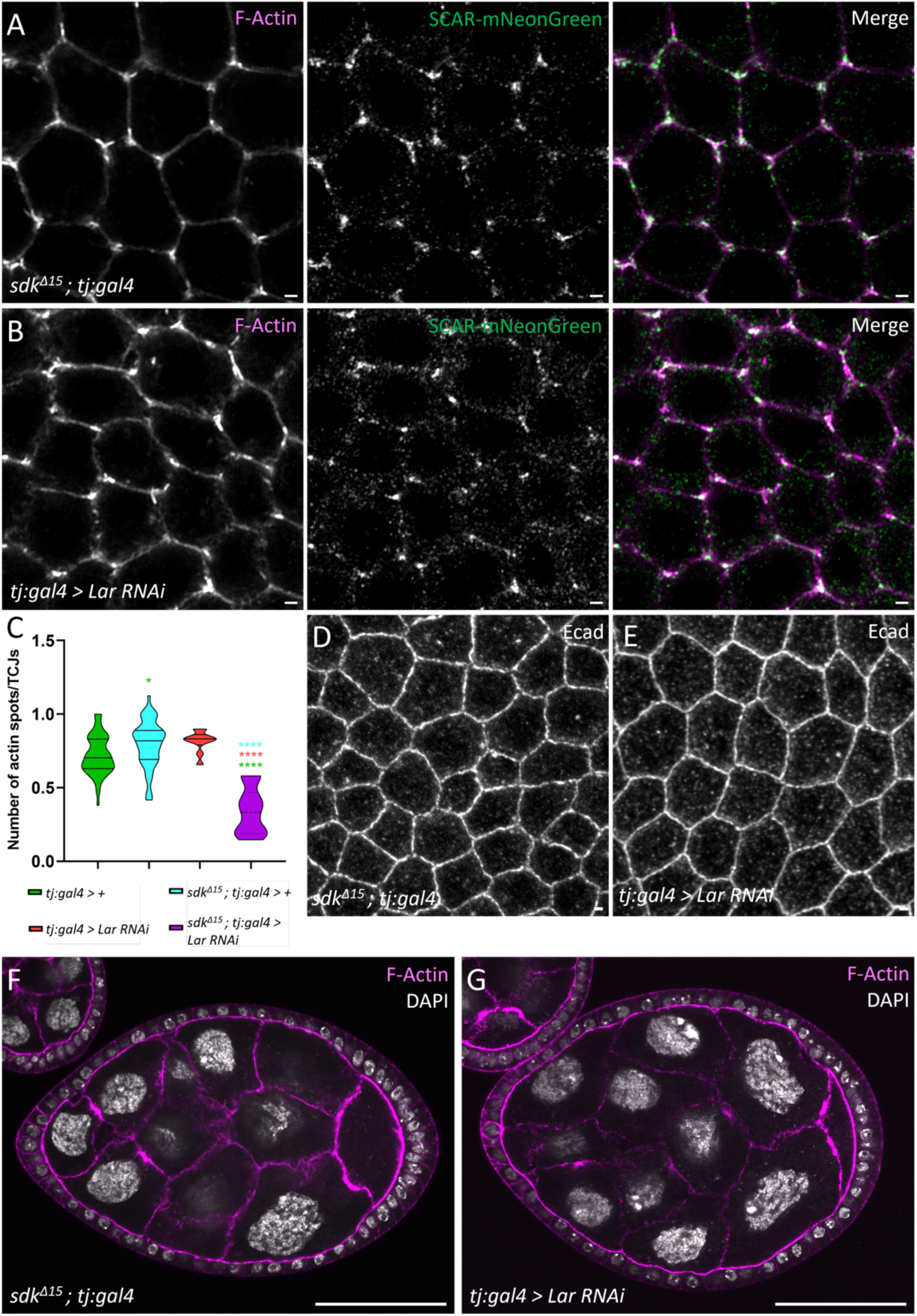
A-B) Scar-mNeonGreen and F-Actin in *sdk* mutant (A) or RNAi against *Lar*. (C) Quantification of F-Actin spots/TCJs in indicated genotypes. (E) *sdk* mutant or (E) RNAi against *Lar* follicles with an immunostaining against Ecad. (F-G) *sdk* mutant and RNAi against *Lar* stage 7 follicles stained with F-Actin and DAPI.

**Table S1.**
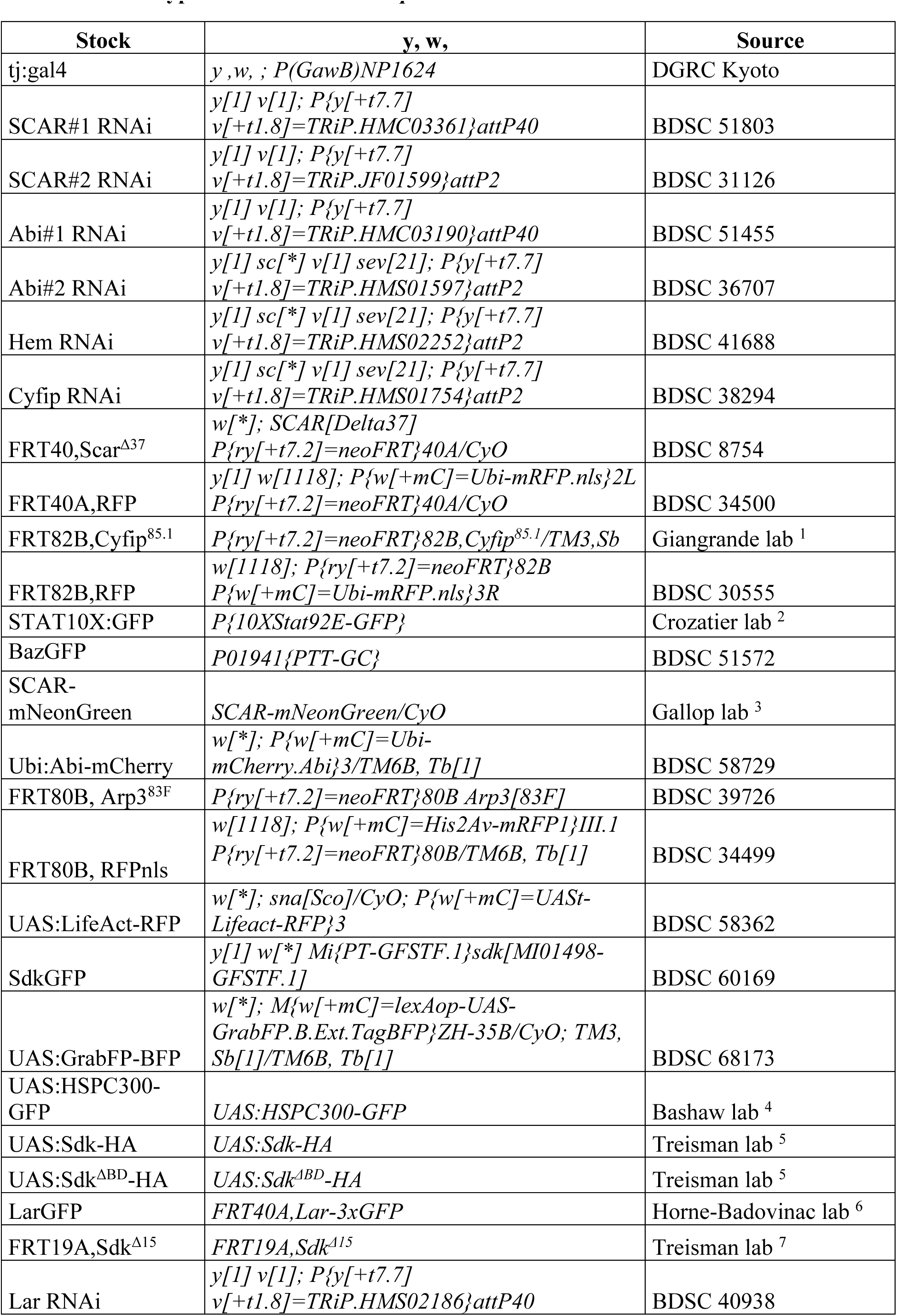
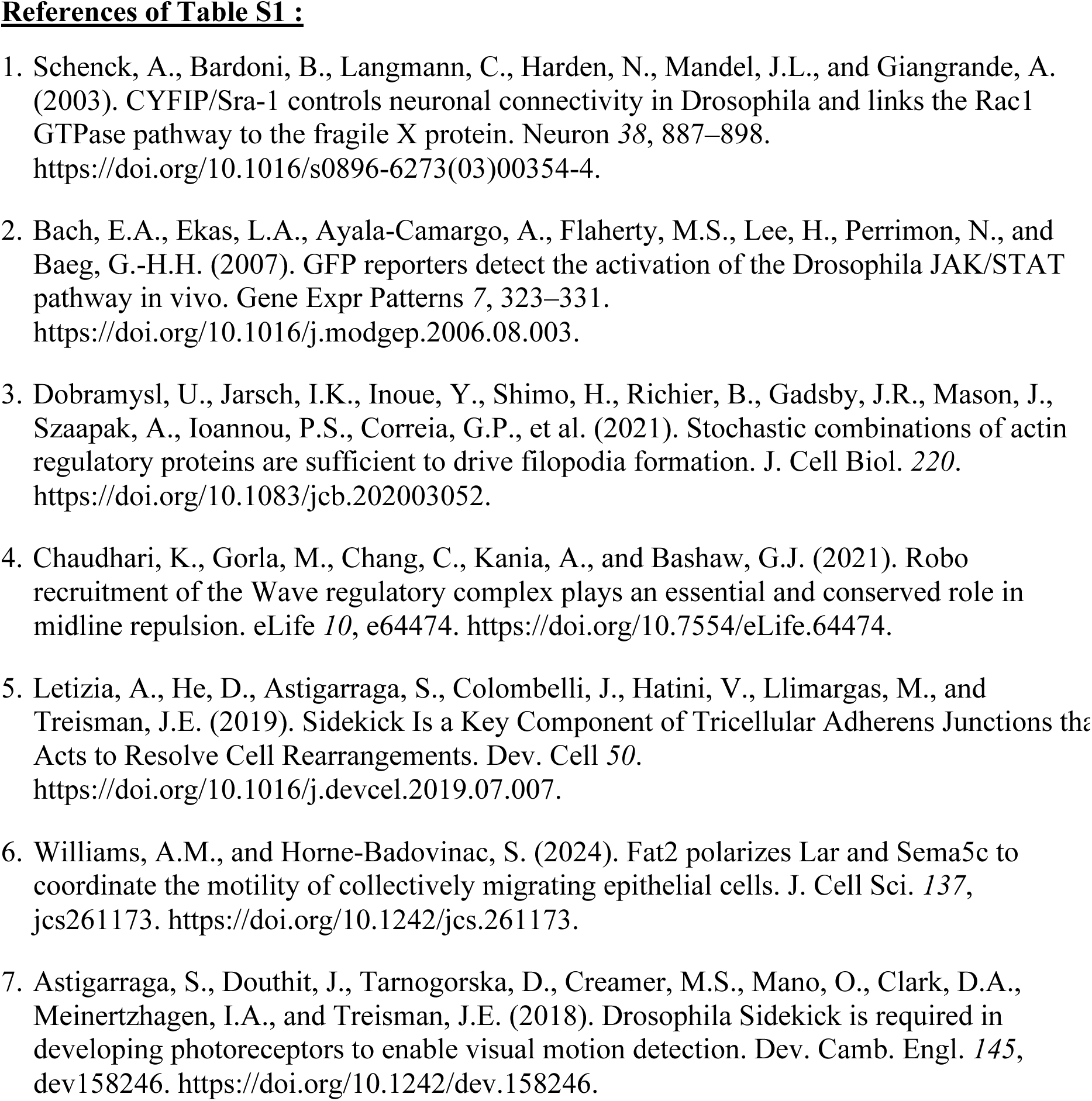
Genotype and source of *Drosophila* strains.

**Table S2.**
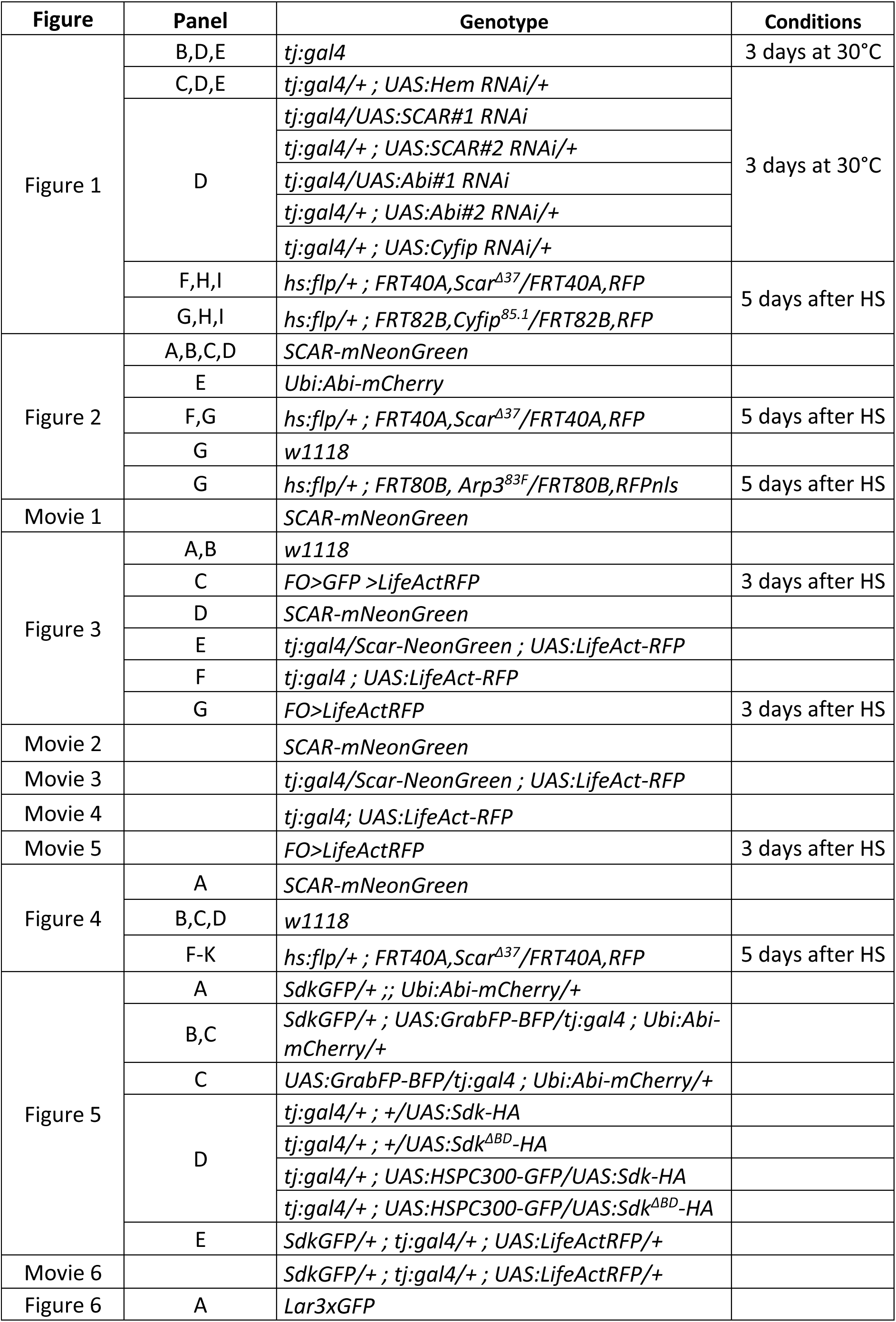
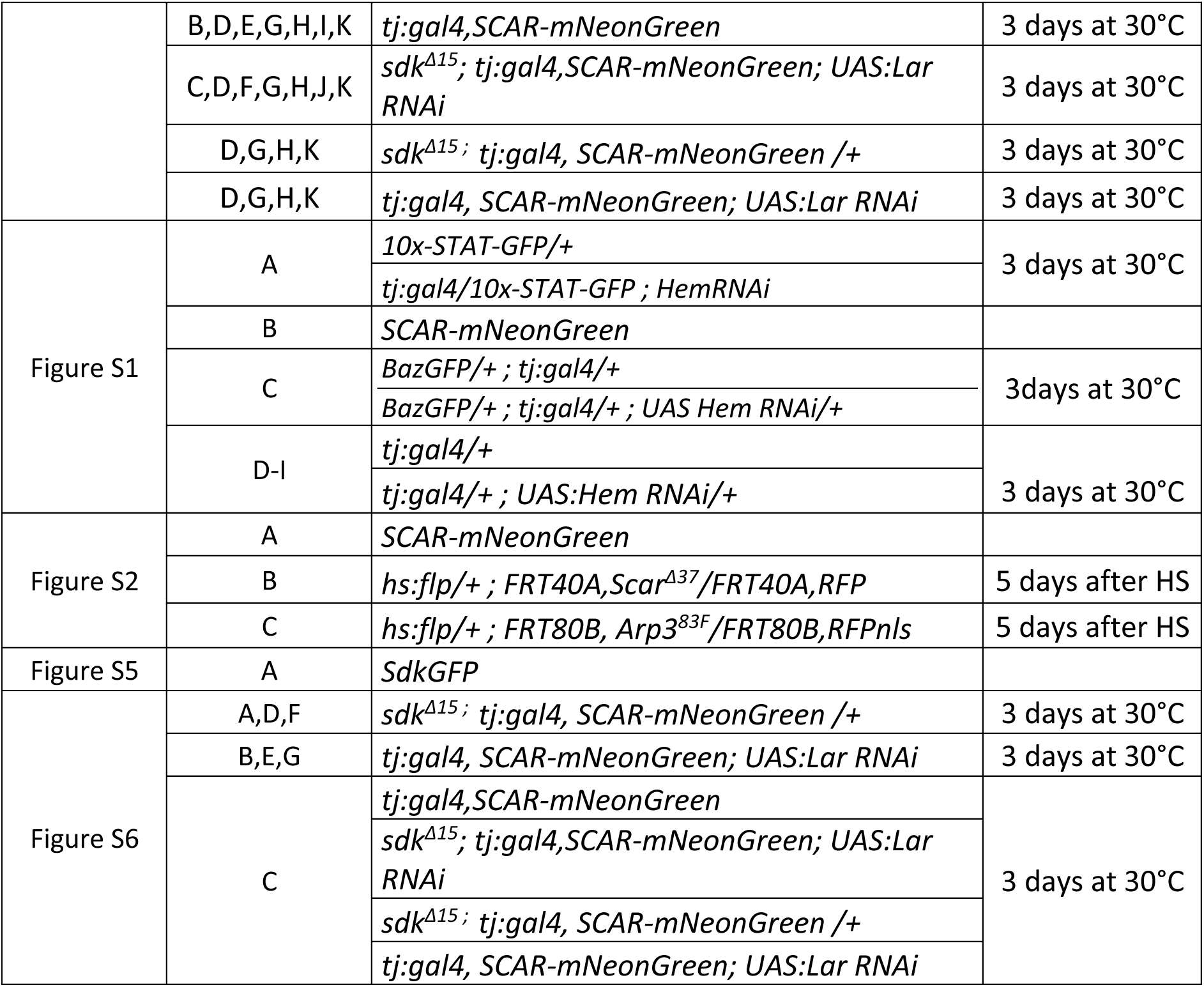
Detailed genotypes and experimental conditions.

| Figure | Panel | Genotype | Conditions |
| --- | --- | --- | --- |
| Figure 1 | B,D,E | *tj:gal4* | 3 days at 30°C |
|  | C,D,E | *tj:gal4/+ ; UAS:Hem RNAi/+* | 3 days at 30°C |
|  | D | *tj:gal4/UAS:SCAR#1 RNAi* |  |
|  |  | *tj:gal4/+ ; UAS:SCAR#2 RNAi/+* |  |
|  |  | *tj:gal4/UAS:Abi#1 RNAi* |  |
|  |  | *tj:gal4/+ ; UAS:Abi#2 RNAi/+* |  |
|  |  | *tj:gal4/+ ; UAS:Cyfp RNAi/+* |  |
|  | F,H,I | *hs:flp/+ ; FRT40A,Scar<sup>Δ37</sup>/FRT40A,RFP* | 5 days after HS |
|  | G,H,I | *hs:flp/+ ; FRT82B,Cyfp<sup>85.1</sup>/FRT82B,RFP* |  |
| Figure 2 | A,B,C,D | *SCAR-mNeonGreen* |  |
|  | E | *Ubi:Abi-mCherry* |  |
|  | F,G | *hs:flp/+ ; FRT40A,Scar<sup>Δ37</sup>/FRT40A,RFP* | 5 days after HS |
|  | G | *w1118* |  |
|  | G | *hs:flp/+ ; FRT80B, Arp3<sup>83F</sup>/FRT80B,RFPnls* | 5 days after HS |
| Movie 1 |  | *SCAR-mNeonGreen* |  |
| Figure 3 | A,B | *w1118* |  |
|  | C | *FO>GFP >LifeActRFP* | 3 days after HS |
|  | D | *SCAR-mNeonGreen* |  |
|  | E | *tj:gal4/Scar-NeonGreen ; UAS:LifeAct-RFP* |  |
|  | F | *tj:gal4 ; UAS:LifeAct-RFP* |  |
|  | G | *FO>LifeActRFP* | 3 days after HS |
| Movie 2 |  | *SCAR-mNeonGreen* |  |
| Movie 3 |  | *tj:gal4/Scar-NeonGreen ; UAS:LifeAct-RFP* |  |
| Movie 4 |  | *tj:gal4; UAS:LifeAct-RFP* |  |
| Movie 5 |  | *FO>LifeActRFP* | 3 days after HS |
| Figure 4 | A | *SCAR-mNeonGreen* |  |
|  | B,C,D | *w1118* |  |
|  | F-K | *hs:flp/+ ; FRT40A,Scar<sup>Δ37</sup>/FRT40A,RFP* | 5 days after HS |
| Figure 5 | A | *SdkGFP/+ ; Ubi:Abi-mCherry/+* |  |
|  | B,C | *SdkGFP/+ ; UAS:GrabFP-BFP/tj:gal4 ; Ubi:Abi-mCherry/+* |  |
|  | C | *UAS:GrabFP-BFP/tj:gal4 ; Ubi:Abi-mCherry/+* |  |
|  | D | *tj:gal4/+ ; +/UAS:Sdk-HA* |  |
|  |  | *tj:gal4/+ ; +/UAS:Sdk<sup>ΔBD</sup>-HA* |  |
|  |  | *tj:gal4/+ ; UAS:HSPC300-GFP/UAS:Sdk-HA* |  |
|  |  | *tj:gal4/+ ; UAS:HSPC300-GFP/UAS:Sdk<sup>ΔBD</sup>-HA* |  |
|  | E | *SdkGFP/+ ; tj:gal4/+ ; UAS:LifeActRFP/+* |  |
| Movie 6 |  | *SdkGFP/+ ; tj:gal4/+ ; UAS:LifeActRFP/+* |  |
| Figure 6 | A | *Lar3xGFP* |  |
|  | B,D,E,G,H,I,K | *tj:gal4,SCAR-mNeonGreen* | 3 days at 30°C |
|  | C,D,F,G,H,J,K | *sdk<sup>Δ15</sup>; tj:gal4,SCAR-mNeonGreen; UAS:Lar RNAi* | 3 days at 30°C |
|  | D,G,H,K | *sdk<sup>Δ15</sup>; tj:gal4, SCAR-mNeonGreen /+* | 3 days at 30°C |
|  | D,G,H,K | *tj:gal4, SCAR-mNeonGreen; UAS:Lar RNAi* | 3 days at 30°C |
| Figure S1 | A | *10x-STAT-GFP/+* | 3 days at 30°C |
|  |  | *tj:gal4/10x-STAT-GFP ; HemRNAi* |  |
|  | B | *SCAR-mNeonGreen* |  |
|  | C | *BazGFP/+ ; tj:gal4/+* | 3days at 30°C |
|  |  | *BazGFP/+ ; tj:gal4/+ ; UAS Hem RNAi/+* |  |
|  | D-I | *tj:gal4/+* | 3 days at 30°C |
|  |  | *tj:gal4/+ ; UAS:Hem RNAi/+* |  |
| Figure S2 | A | *SCAR-mNeonGreen* |  |
|  | B | *hs:flp/+ ; FRT40A,Scar<sup>Δ37</sup>/FRT40A,RFP* | 5 days after HS |
|  | C | *hs:flp/+ ; FRT80B, Arp3<sup>83F</sup>/FRT80B,RFPnls* | 5 days after HS |
| Figure S5 | A | *SdkGFP* |  |
| Figure S6 | A,D,F | *sdk<sup>Δ15</sup>; tj:gal4, SCAR-mNeonGreen /+* | 3 days at 30°C |
|  | B,E,G | *tj:gal4, SCAR-mNeonGreen; UAS:Lar RNAi* | 3 days at 30°C |
|  | C | *tj:gal4,SCAR-mNeonGreen* | 3 days at 30°C |
|  |  | *sdk<sup>Δ15</sup>; tj:gal4,SCAR-mNeonGreen; UAS:Lar RNAi* |  |
|  |  | *sdk<sup>Δ15</sup>; tj:gal4, SCAR-mNeonGreen /+* |  |
|  |  | *tj:gal4, SCAR-mNeonGreen; UAS:Lar RNAi* |  |

**Table S3.**
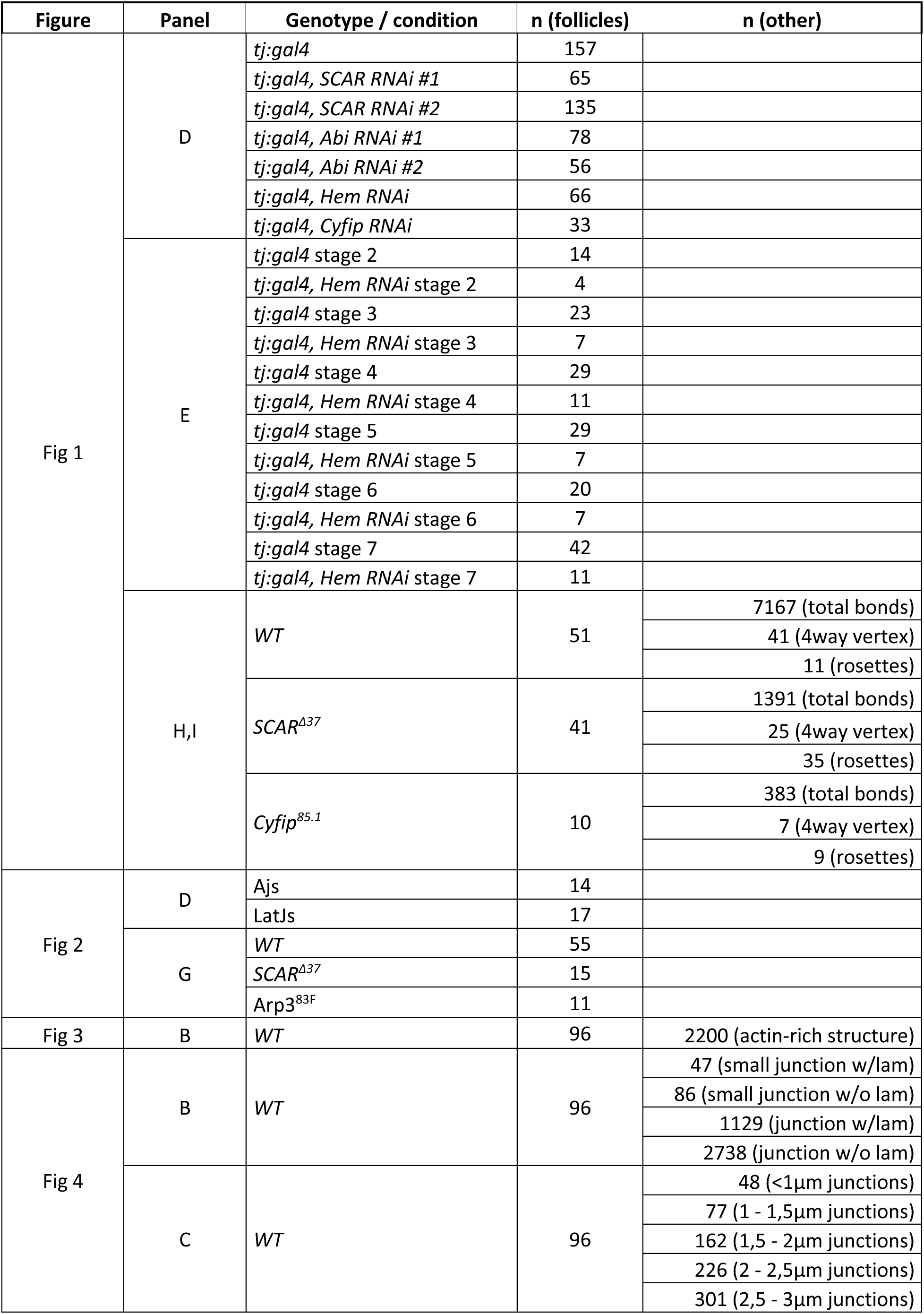
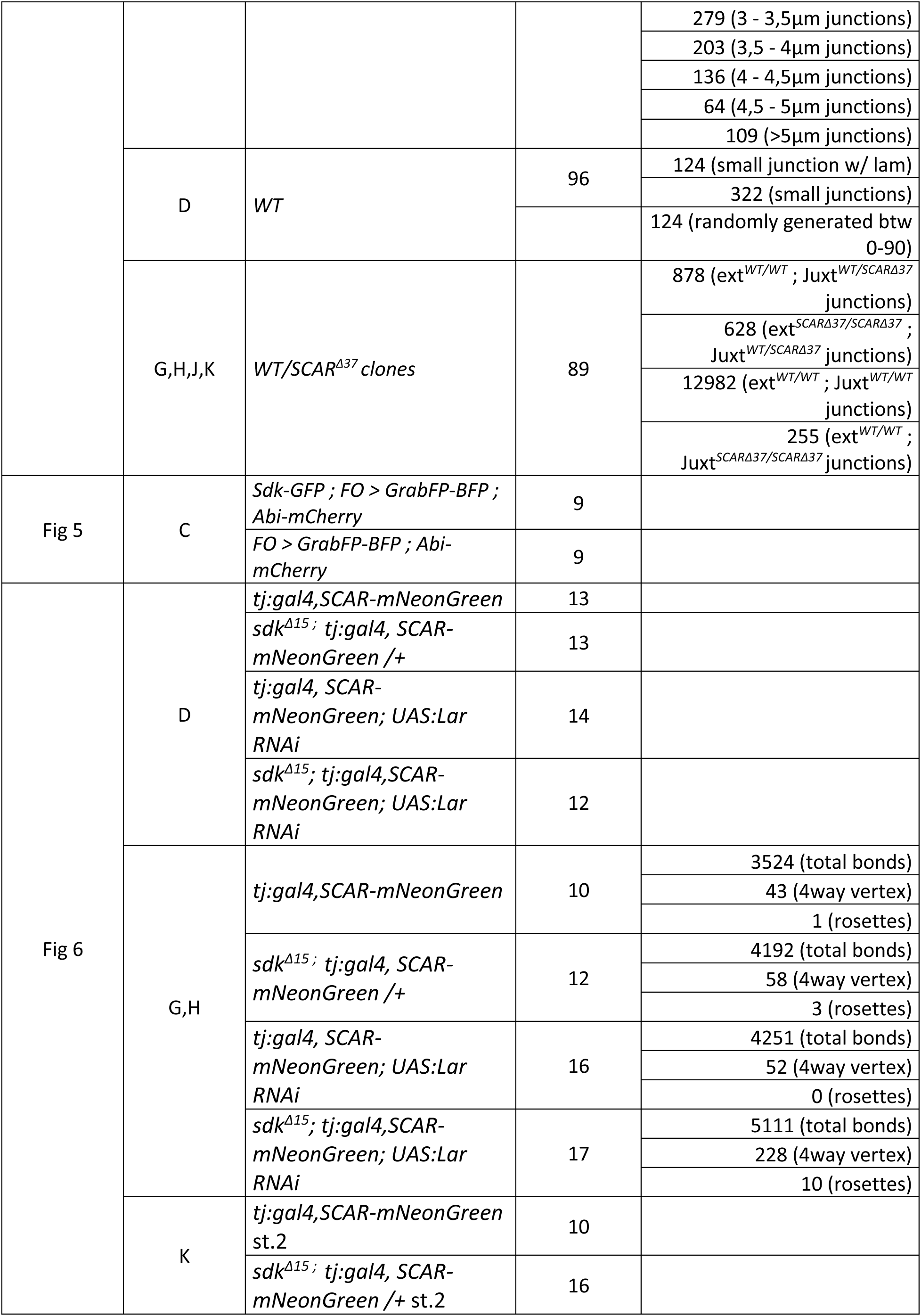
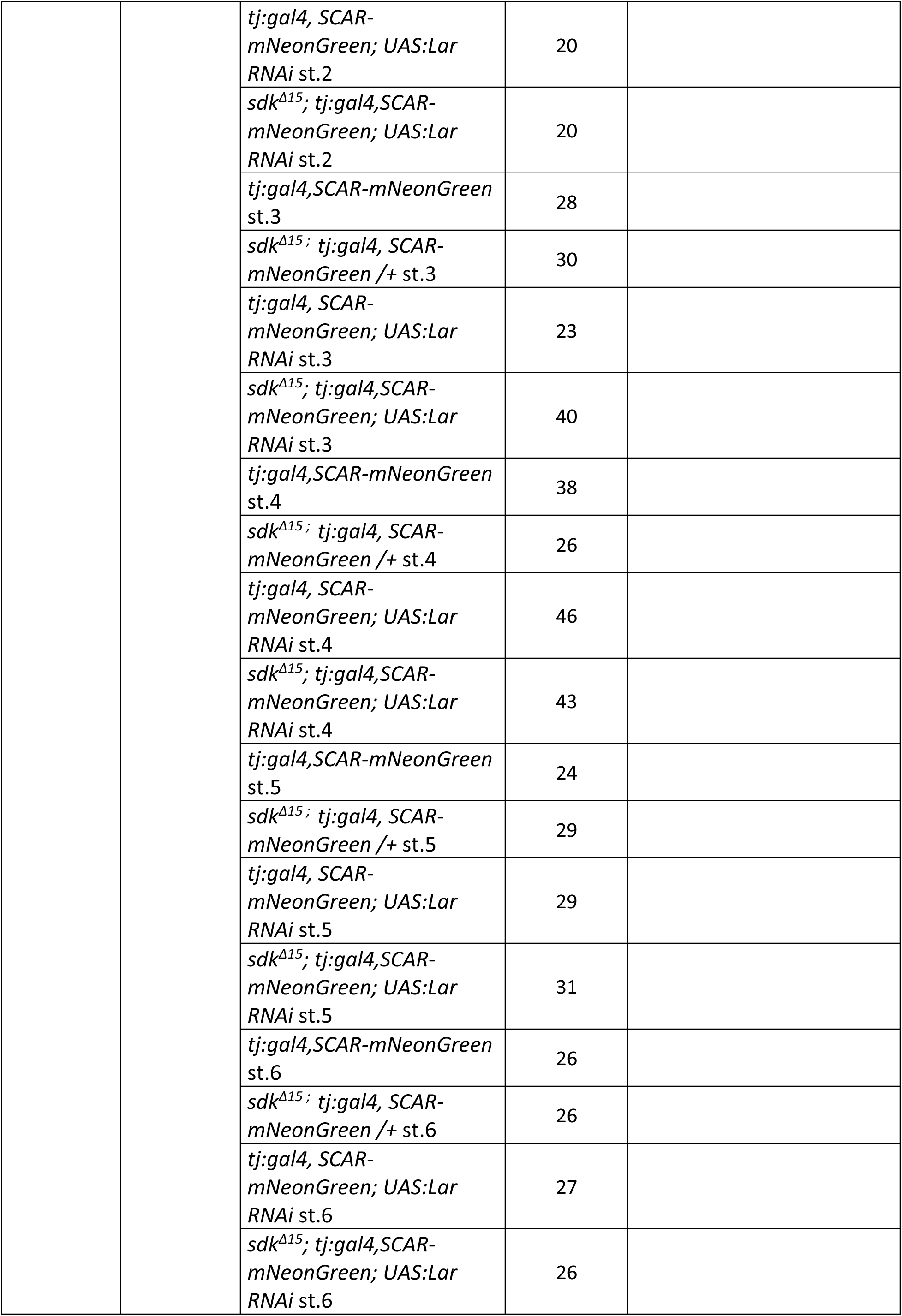
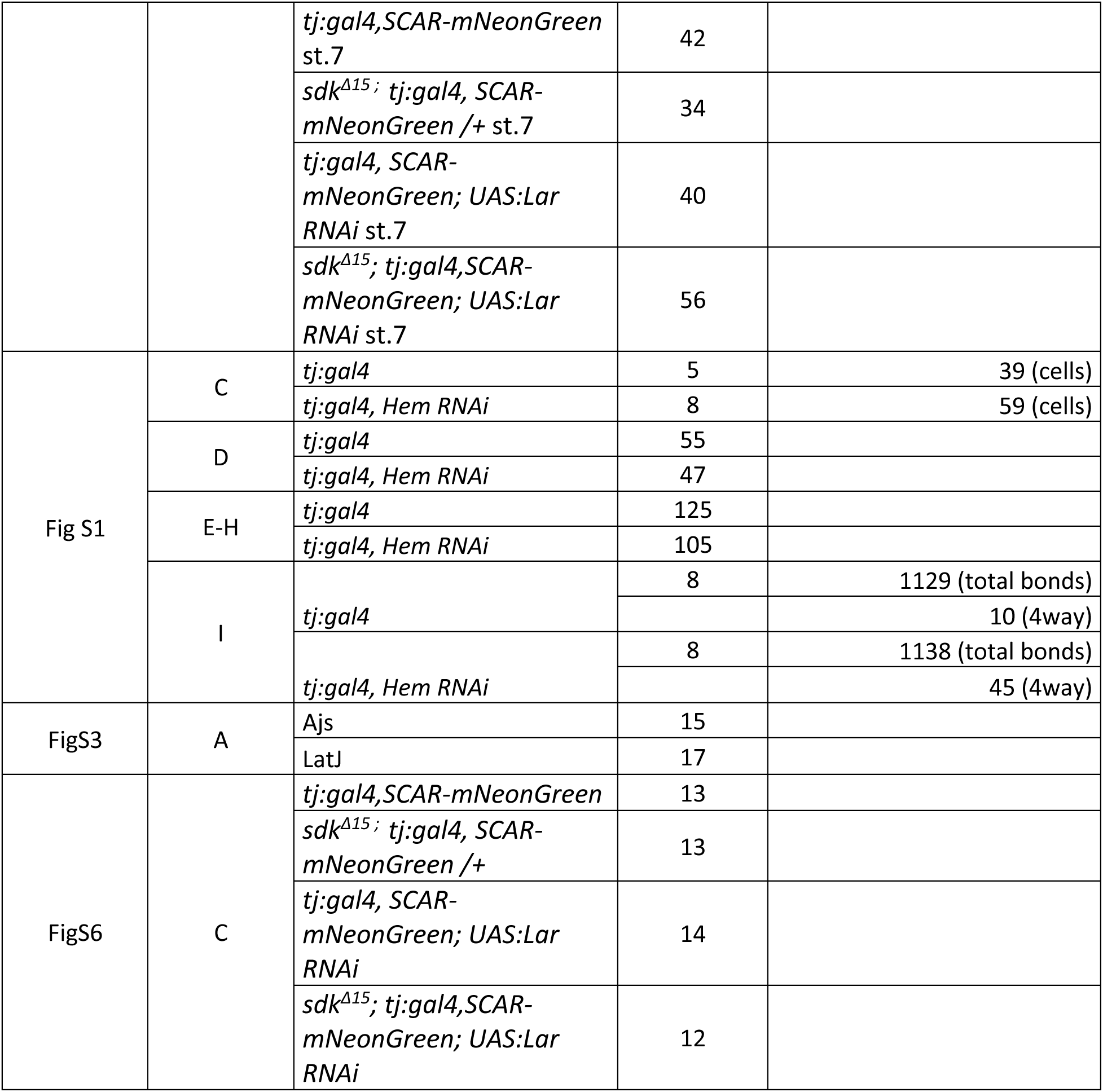
Detailed sample size.

